# The BLM-TOP3A-RMI1-RMI2 proximity map reveals that RAD54L2 suppresses sister chromatid exchanges

**DOI:** 10.1101/2024.04.07.588476

**Authors:** Jung Jennifer Ho, Edith Cheng, Cassandra J. Wong, Jonathan R. St-Germain, Wade H. Dunham, Brian Raught, Anne-Claude Gingras, Grant W. Brown

## Abstract

Homologous recombination is a largely error-free DNA repair mechanism conserved across all domains of life and is essential for the maintenance of genome integrity. Not only are the mutations in homologous recombination repair genes probable cancer drivers, some also cause genetic disorders. In particular, mutations in the Bloom (BLM) helicase cause Bloom Syndrome, a rare autosomal recessive disorder characterized by increased sister chromatid exchanges and predisposition to a variety of cancers. The pathology of Bloom Syndrome stems from the impaired activity of the BLM-TOP3A-RMI1-RMI2 (BTRR) complex which suppresses crossover recombination to prevent potentially deleterious genome rearrangements. We provide a comprehensive BTRR proximity interactome, revealing proteins that suppress crossover recombination. We find that RAD54L2, a SNF2-family protein, physically interacts with BLM and suppresses sister chromatid exchanges. RAD54L2 is important for recruitment of BLM to chromatin and requires an intact ATPase domain to promote non-crossover recombination. Thus, the BTRR proximity map identifies a regulator of recombination.

## Introduction

The repair of double-strand DNA breaks (DSBs) by homologous recombination (HR) is critical for the maintenance of genome stability. HR is a high-fidelity repair process that employs homologous DNA sequences, typically in the sister chromatid, as a template to repair DSBs while retaining critical sequence information. Mutations in HR genes can cause genome instability and predisposition to cancer; for example, ataxia telangiectasia, Nijmegen breakage syndrome, and Bloom syndrome are caused by mutations in ATM, NBN, and BLM, respectively, and are associated with pleiotropic cancer susceptibility (Renwick *et al*, 2006; Belhadj *et al*, 2023; German, 1997). Bloom syndrome is a rare disease characterized by small stature, extreme sensitivity to sunlight, immunodeficiency, and oncogenesis (reviewed in (de Renty & Ellis, 2017)). The DNA helicase BLM plays critical roles in DSB repair to suppress oncogenesis. BLM functions in a complex with the topoisomerase TOP3A (Hu *et al*, 2001; Wu *et al*, 2000) and the structural components RMI1 (Meetei *et al*, 2003; Yin *et al*, 2005; Raynard *et al*, 2006) and RMI2 (Singh *et al*, 2008) to suppress rearrangements during HR repair of DSBs (Bythell-Douglas & Deans, 2021) and stressed DNA replication forks (Lönn *et al*, 1990; Sengupta *et al*, 2003; Davies *et al*, 2004, 2007), thereby maintaining genome integrity.

BLM responds to the presence of DSBs both early and late in HR repair. In the early stages of repair, BLM DNA helicase activity promotes the long-range resection of double-stranded DNA ends in concert with the DNA2 endonuclease (Gravel *et al*, 2008; Nimonkar *et al*, 2011). The resulting 3’ single-stranded DNA tails are substrates for the assembly of RAD51-ssDNA filaments, which conduct homology searches for suitable repair templates (Zhao *et al*, 2015). In the late stages of HR repair, BLM functions in a complex with TOP3A (Hu *et al*, 2001; Wu *et al*, 2000), RMI1 (Yin *et al*, 2005; Wu *et al*, 2006), and RMI2 (Singh *et al*, 2008) to dissolve four-way recombination intermediates known as double Holliday junctions (dHJs) (Bizard & Hickson, 2014). Dissolution of dHJs by the BLM-TOP3A-RMI1-RMI2 (BTRR) complex is distinguished from dHJ resolution by structure-selective nucleases in that dissolution produces exclusively non-crossover products. As such, defects in BTRR components result in the canonical phenotype of BLM syndrome: increased sister chromatid exchanges (SCEs) (German *et al*, 1977; Martin *et al*, 2018; Gönenc *et al*, 2022; Hudson *et al*, 2016), increased loss of heterozygosity events (Yusa *et al*, 2004; Langlois *et al*, 1989; LaRocque *et al*, 2011), and increased risks of pleiotropic cancer (German, 1997; Ababou, 2021). Elevated SCEs are a broad indicator of genome instability and occur at random locations across the mammalian genome (Hamadeh et al., 2022) and in particular, in hotspots at common fragile sites (van Wietmarschen *et al*, 2018). SCEs are associated with the use of therapeutics that stall DNA replication such as PARP inhibitors (Heijink *et al*, 2022), and those that generate DSBs such as irradiation (Conrad *et al*, 2011).

In addition to its primary function in the suppression of crossover recombination, BLM functions in many other aspects of genome stability. These include the restart of stalled replication forks (Davies *et al*, 2007), unwinding of RNA G-quadruplexes in stress granules (Danino *et al*, 2023), unwinding of DNA G-quadruplexes during telomere replication (Drosopoulos *et al*, 2015), and the repair of ultra-fine DNA bridges in anaphase (Chan *et al*, 2007). While there are no separation-of-function mutations that define each role of BLM in the maintenance of genome stability, it is thought that BLM function is largely defined by its interacting partners and nucleic acid substrates.

BLM-interacting proteins have been identified by several affinity-purification/mass spectrometry approaches, including affinity purification of streptavidin-tagged BLM (Wan *et al*, 2013; Wang *et al*, 2013), immunoprecipitation of endogenous BLM using anti-BLM antibodies (Yin *et al*, 2005; Meetei *et al*, 2003; Guo *et al*, 2023; Bhattacharyya *et al*, 2009), and affinity purification of engineered fluorescent protein BLM fusions (Hein *et al*, 2015; Cho *et al*, 2022), each coupled to mass spectrometry to identify interacting proteins. Despite these comprehensive affinity-purification interactomes, BTRR is often found in nuclear foci (Wang *et al*, 2022; Eladad *et al*, 2005; Davalos *et al*, 2004),which are biochemically unstable and insoluble (Takata *et al*, 2009), making it challenging to capture a complete and physiologically relevant BTRR interactome.

Here, we map the BTRR interactome using proximity-dependent biotin identification (BioID) and identify proteins that suppress crossover recombination. We define 206 high-confidence BTRR proximity interactions, including 24 proximity interactions that are shared by at least two BTRR subunits. Comparison of N-versus C-terminal BLM BioID suggests most BLM interactions occur in proximity to the N-terminus. Analysis of the BTRR interactome revealed 15 proteins that suppress SCEs, including the SNF2-family chromatin remodeler RAD54L2. Knockout of *RAD54L2* results in increased SCEs and decreased non-crossover recombination. RAD54L2 is important for recruitment of BLM to repair foci, but does not influence MRE11 or RAD51 recruitment, indicating that RAD54L2 likely functions during dissolution of dHJs. Thus, we identify a new player in the processing of recombination intermediates.

## Results

### The BTRR proximity interactome

To capture the proximity interactome of the BTRR complex, we performed BioID to selectively biotinylate proteins in close proximity to each BTRR complex member (Roux *et al*, 2012). We generated four tetracycline-inducible HEK293 cell lines that stably expressed N-terminal BirA* tagged BTRR fusion proteins (Fig EV1). Following 24-hour induction of each BirA* fusion protein, biotinylated proteins were affinity-purified and analysed by mass spectrometry. The confidence of each BTRR proximity interaction was assessed with SAINTexpress (Teo *et al*, 2014) to predict the likelihood of a true interaction for each bait-prey pair. We detected a total of 64 high-confidence proximal proteins (Bayesian false discovery rate (BFDR) ≤ 0.01) for BLM, 18 for TOP3A, 84 for RMI1, and 40 for RMI2 (Fig 1A,B).

**Figure 1.**
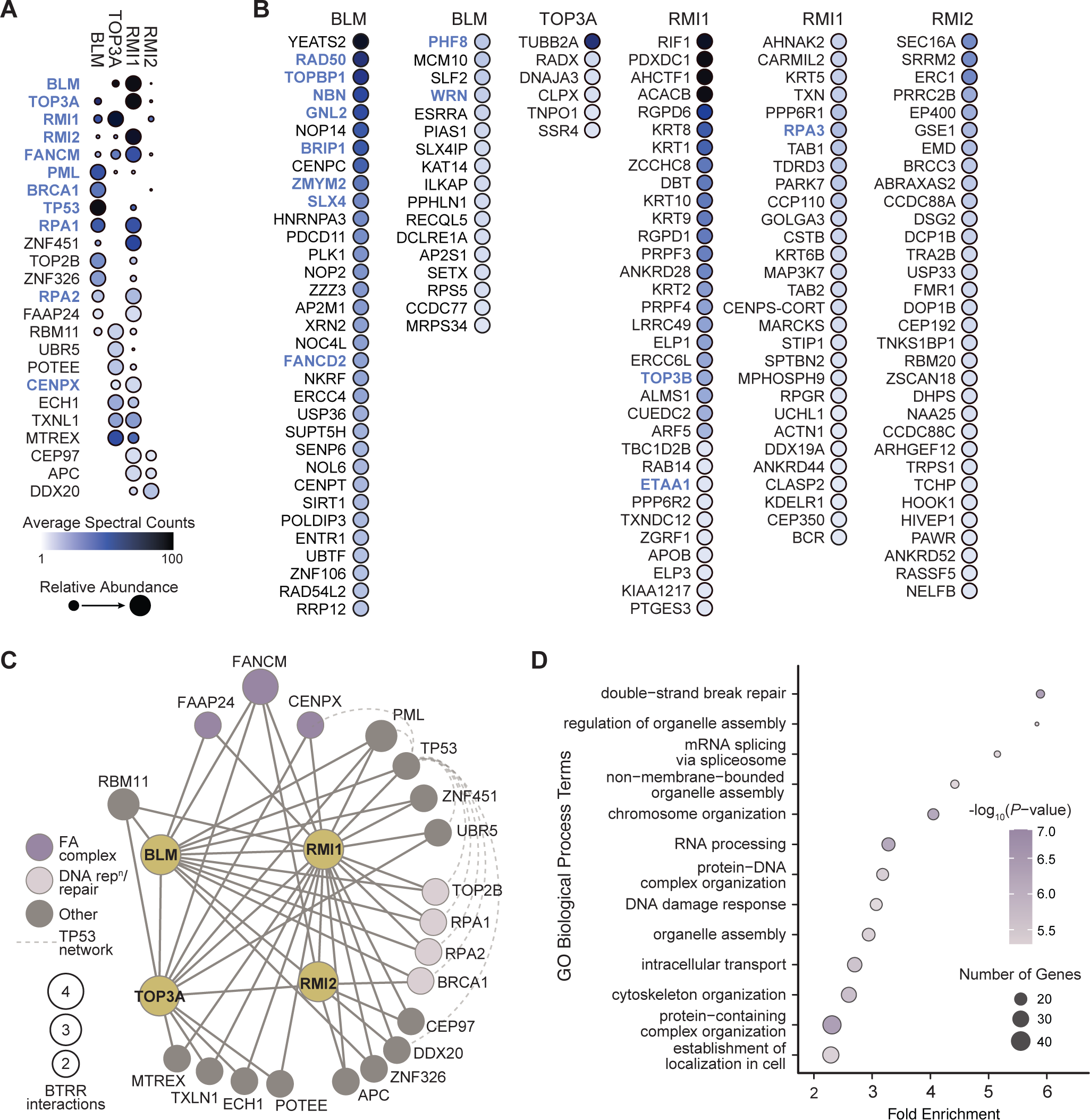
BioID identifies proximal interactors of BLM, TOP3A, RMI1, and RMI2. A. Dot plot showing prey proteins identified with BirA*-BLM, BirA*-TOP3A, BirA*-RMI1, and BirA*-RMI2. High-confidence proximal proteins identified with at least two baits are shown (BFDR ≤ 0.01). BioID screens were performed in duplicate (BLM and TOP3A), triplicate (RMI2), or sextuplicate (RMI1). Dot colour indicates the average spectral counts, and dot size indicates relative abundance across the baits. Previously described interaction partners annotated in BioGRID are shown in blue. B. Dot plot showing all high-confidence prey proteins identified with one BTRR bait, with BFDR ≤ 0.01. Details as in panel A. C. Network of the 24 proximal proteins detected in at least two BTRR BioID screens. The nodes identify prey proteins and are sized according to the number of proximity interactions. The solid edges connect the BioID BTRR baits with their proximal preys. The dashed edges indicate protein-protein interactions among a subset of the BTRR proximal proteins as annotated in BioGRID. Members of the Fanconi Anemia (FA) core complex, DNA replication/repair proteins, and a TP53 interaction network are indicated. D. Gene ontology (GO) biological process analysis for 150 high-confidence BTRR proximal proteins not currently annotated in BioGRID. The-fold enrichment for each GO term is indicated, the colours indicate the corrected P-values, and the size of the circles corresponds to the number of genes annotated to the given GO term.

Previous analyses have detected diverse proteins physically interacting, directly or indirectly, with BTRR. Comparing to protein interactors with low-throughput evidence in the BioGRID database (Oughtred *et al*, 2021) (https://thebiogrid.org/; accessed 26/09/2023; Source Data 1A,B), we detected 15 of 78 annotated interactions for BLM, 3 of 18 for TOP3A, 6 of 10 for RMI1, and 3 of 8 for RMI2 (Fig 1A,B and Source Data 1A,B). Notably, the known interacting proteins of BTRR that we detected with BioID include components of the Fanconi Anemia (FA) complex (FANCM, FANCD2, BRIP1/FANCJ, and CENPX) involved in double-strand break repair and DNA replication stress response (Meetei *et al*, 2003), double-strand break repair proteins (RAD50 (Franchitto & Pichierri, 2002) and BRCA1 (Acharya *et al*, 2014)), and proteins that repair stalled replication forks (TOPBP1 (Wang *et al*, 2013), ETAA1 (Bass *et al*, 2016), WRN (Sturzenegger *et al*, 2014)). Thus, the proximity interactome of BTRR contains the expected network of DNA damage and replication stress response and repair proteins.

Twenty-four proximal proteins were detected in at least two BTRR BioID screens (Fig 1C and Source Data 1C), 14 of which are not currently annotated as BTRR interactors in BioGRID. The known BTRR binding partner FANCM (Lu *et al*, 2019) was identified in all 4 screens. Interaction with FAAP24, an additional member of the FA complex (Coulthard *et al*, 2013) not currently known to interact with BTRR, was identified in 3 screens, as was the FA complex interactor CENPX. Analysis in GeneMANIA (Franz *et al*, 2018) (https://genemania.org/; accessed 24/01/2024) revealed an interacting network of proteins that were identified in at least 2 screens, including the tumour suppressor TP53, the p53 regulator PML, the DNA damage responsive E3 ligases UBR5 and ZNF451, and the DNA repair proteins BRCA1, CENPX, TOP2B, RPA1, and RPA2 (Fig 1C and Source Data 1C).

We identified a total of 167 interactions that are not currently annotated in either low-throughput or high-throughput datasets (Oughtred *et al*, 2021). We performed a gene ontology analysis of the corresponding 150 genes, discovering statistically supported enrichments for DNA repair, DNA damage response, chromosome organization genes, and an unexpected enrichment for genes involved in non-membrane-bounded organelle assembly particularly genes involved in kinetochore and centrosome function (Fig 1D and Source Data1D).

### The BLM N- and C-terminus proximity interactome

Since BLM is a 159 kDa multi-domain protein that binds protein partners via both N- and C-terminal regions (Yin *et al*, 2005; Wan *et al*, 2013; Meetei *et al*, 2003; Guo *et al*, 2023; Bhattacharyya *et al*, 2009), we aimed to define whether fusing a biotin ligase at the N- or C-terminus would reveal different proximal interactomes. We therefore fused the biotin ligase miniTurbo (mT) (Branon *et al*, 2018) to the N- or the C-terminus of BLM (Fig 2). SAINTexpress (Teo *et al*, 2014) analysis of the miniTurbo proximity labeling data revealed 51 high-confidence proximity interactions with the N-terminal BLM fusion (mT-BLM) and 15 interactions with the C-terminal BLM fusion (BLM-mT) with BFDR ≤ 0.01 (Figure 2A, Source Data 2A). Most of the interactions were detected with the N-terminal fusion, consistent with most of the annotated BLM interactions occurring via the amino terminus (Yin *et al*, 2005; Wan *et al*, 2013; Meetei *et al*, 2003; Guo *et al*, 2023; Bhattacharyya *et al*, 2009; Yankiwski *et al*, 2001). One exception was PML, which was detected with BirA*-TOP3A, BirA*-RMI1, and BLM-mT, suggesting that PML is not in close proximity to the BLM amino terminus. BLM localization to PML nuclear bodies appears to be dependent on the amino terminus (Yankiwski *et al*, 2001), indicating that localization to PML bodies and proximity to PML are not necessarily concordant.

**Figure 2.**
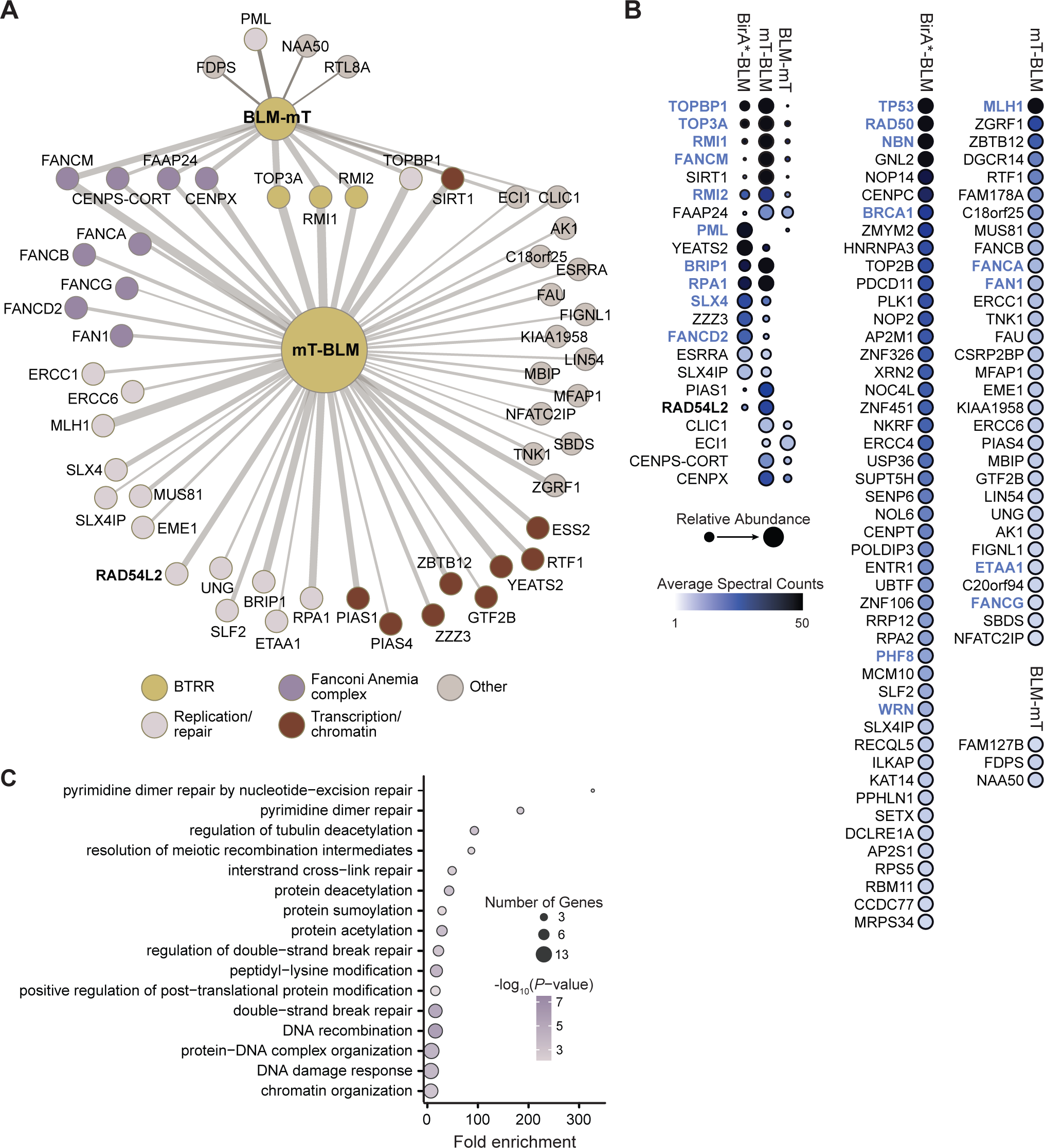
High-resolution proximal interactome of the BLM protein. A. Network of the 15 proximal proteins detected with BLM tagged with miniTurbo at the C-terminus (BLM-mT) and the 51 proximal proteins detected with BLM tagged at the N-terminus (mT-BLM). The nodes are the proximal protein preys and the edge widths are proportional to the log2 of the average spectral counts for the given bait-prey pair. Members of the Fanconi Anemia complex, the BTRR complex, proteins that function in DNA replication and/or DNA repair, and proteins that function in transcription and/or chromatin function are indicated. B. Dot plot showing prey proteins identified with BLM N- and C-terminal miniTurbo fusions. High-confidence proximal proteins are shown (BFDR ≤ 0.01), with the preys from the BirA*-BLM screen from Fig. 1A plotted for comparison. Each BioID screen was performed in quadruplicate, with dot colour indicating the average spectral counts. The dot outlines indicate the BFDR, and dot size indicates relative abundance across the baits. Previously described interaction partners annotated in BioGRID are shown in blue. C. Gene ontology (GO) biological process analysis for 101 high-confidence BLM proximal proteins identified in the BirA* and miniTurbo BioID screens. The-fold enrichment for each GO term is indicated, the colours indicate the corrected P-values, and the size of the circles corresponds to the number of genes annotated to the given GO term.

We next combined the data from the three BLM BioID screens (BirA*-BLM, mT-BLM, and BLM-mT; Fig 2B, Source Data 1AB, and Source Data 2B). The BLM proximity interactome comprises 101 high confidence proximal proteins, and is highly enriched for annotated BLM protein-protein interactions (Oughtred *et al*, 2021) (27-fold; hypergeometric p = 1.3 x 10^-30^). Of note, we identified 76 interactions that are not currently annotated, expanding the BLM interactome to 259 proteins. Functional analysis of the BLM proximity interactome (Fig 2C and Source Data 2C) indicated the expected enrichment for DNA recombination and repair, and revealed extensive interactions with post-translational modification pathways including sumoylation and acetylation.

### Proximal partners of BTRR suppress sister chromatid exchanges

To identify the high confidence proximity interactions that are most relevant to BTRR function, we selected a subset of proximal proteins from our BTRR BioID screens for sister chromatid exchange (SCE) analysis. An increase in SCEs is the hallmark cellular phenotype of Bloom syndrome cells (German *et al*, 1977), and increased SCEs are evident when any component of BTRR is depleted (Martin *et al*, 2018; Hudson *et al*, 2016; Hoadley *et al*, 2010). We selected 18 genes from the BTRR proximity interactome to knock down with siRNA, and included *BLM*, *TOP3A*, and *RMI1* as positive controls. Following siRNA depletion of each gene transcript, we stained the sister chromatids and prepared metaphase chromosome spreads (Fig 3A). We assessed the effect of siRNA depletion of each candidate interactor on the number of SCEs per mitosis (Fig 3B and Source Data 3B). As expected, U2OS cells depleted of BLM or its known interacting partner, RMI1 showed increased frequency of SCEs. Increased SCEs were evident for many candidate genes, encouraging more detailed analyses. In particular, 15 of the 18 gene knockdowns resulted in a statistically supported increase in SCEs, 13 of which showed a greater than 2-fold increase (Fig 3B). We performed additional replicates to validate three candidates that exhibited high frequencies of SCEs (Fig 3C and EV3A). Knockdown of *RAD54L2*, *ABRAXAS2*, or *ZZZ3* using siRNA caused a statistically supported increase in SCEs. The reciprocal of an SCE is non-crossover (NCO) recombination. We used a direct-repeat recombination assay (Weinstock *et al*, 2006), where NCOs reconstitute GFP after a DSB is induced, to assess NCO recombination following siRNA knockdown of *RAD54L2*, *ABRAXAS2*, or *ZZZ3* (Fig 3D and EV3B). Knockdown of each of the three genes resulted in decreased NCO recombination, consistent with the increase in SCEs we found upon knockdown. We infer that *RAD54L2*, *ABRAXAS2*, and *ZZZ3* are important for suppressing crossovers following DSBs.

**Figure 3.**
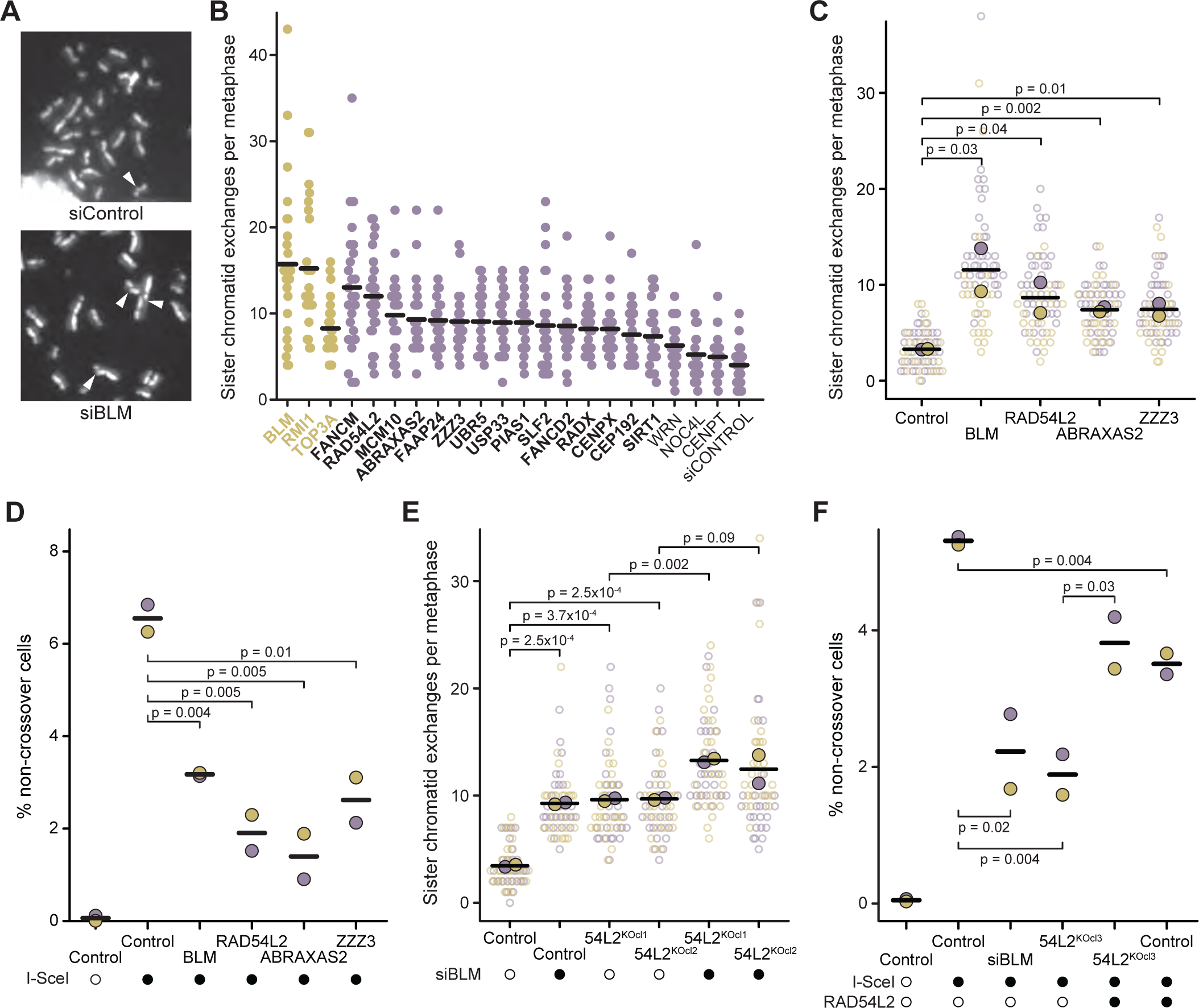
BTRR proximal proteins suppress sister chromatid exchanges. A. Mitotic chromosome spreads from U2OS cells following siRNA knockdown of BLM or control. Sister chromatids are differentially stained to detect exchanges, indicated by white arrows. B. The number of SCEs for 25 metaphases is plotted for the indicated gene knockdowns. Black bars indicate the means. Knockdowns that resulted in a statistically supported increase in SCEs are in bold (p < 0.05, Wilcoxon rank sum test). C. The number of SCEs per metaphase is plotted for two independent replicates of each of the indicated gene knockdowns. The replicates are indicated by the different colours and the mean SCEs for each replicate is indicated by the filled circles. The p-values from one-sided Student’s t-tests comparing the replicate means are indicated. D. The percent of cells expressing GFP (% non-crossover cells) is plotted for two independent replicates of each of the indicated gene knockdowns. Closed circles indicate experiments where a double-strand DNA break was induced by expression of I-SceI. The p-values from one-sided Student’s t-tests are indicated. E. The number of SCEs per metaphase is plotted for two independent replicates for each of two clonal RAD54L2 gene disruption lines. BLM was knocked down with siRNA where indicated (closed circles). The replicates are indicated by the different colours and the mean SCEs for each replicate is indicated by the filled circles. The p-values from one-sided Student’s t-tests comparing the replicate means are shown. F. The percent of cells expressing GFP (% non-crossover cells) is plotted for a clonal RAD54L2 gene disruption line. Where indicated, the wildtype RAD54L2 gene was introduced into the knockout line by transient transfection (closed circles). BLM was knocked down with siRNA as a positive control. A double-strand DNA break was induced by expression of I-SceI where indicated (closed circles). The p-values from one-sided Student’s t-tests are shown.

### *RAD54L2* knockout increases crossover recombination

To confirm the role of RAD54L2 in suppressing crossovers we used CRISPR/Cas9 to disrupt *RAD54L2* in U2OS cells. We tested two independent gene disruption lines for the frequency of SCEs (Fig 3E and EV4A). In both cases, the number of SCEs per mitosis increased when RAD54L2 was disrupted. To gain additional insight into the function of RAD54L2 relative to that of BLM, we knocked down *BLM* in the *RAD54L2* deficient lines and measured SCEs (Fig 3E and EV4B). In both *RAD54L2* deficient lines, knockdown of *BLM* increased the number of SCEs. However, the increase in SCEs was neither multiplicative nor additive, consistent with *BLM* and *RAD54L2* functioning in the same genetic pathway to suppress crossover events, perhaps in addition to making independent contributions to SCE suppression.

We tested the role of RAD54L2 in noncrossover recombination by disrupting *RAD54L2* in the NCO assay cell line (Fig 3F and EV4C). Disruption of *RAD54L2* resulted in a decrease in NCO events. We introduced a wild-type copy of *RAD54L2* into the line, which resulted in a partial rescue of NCO events (Fig 3F, EV4D, and EV4E). Interestingly, expression of wild type *RAD54L2* in the control line caused some decrease in NCO events, suggesting that excess *RAD54L2* might reduce NCO recombination.

### RAD54L2 is proximal to DNA repair proteins and chromatin regulators

To complement our mass spectrometry data, we tested whether we could detect RAD54L2 among the proteins in miniTurbo BLM (Fig 4A) and BLM-BirA* cell lysates (Apppendix Fig S1A). RAD54L2 was detected in streptavidin precipitates only when miniTurbo BLM or BLM-BirA* were expressed, confirming the identification of RAD54L2 in BLM BioID experiments. We then asked if BLM immunoprecipitates contain RAD54L2 (Fig 4B). We immunoprecipitated BLM-BirA* from nuclear extracts with antibodies against the FLAG epitope and probed immunoblots for BLM and RAD54L2. RAD54L2 was specifically present in the immunoprecipitates when BLM-BirA* was expressed, indicating that RAD54L2 and BLM form a stable complex.

**Figure 4.**
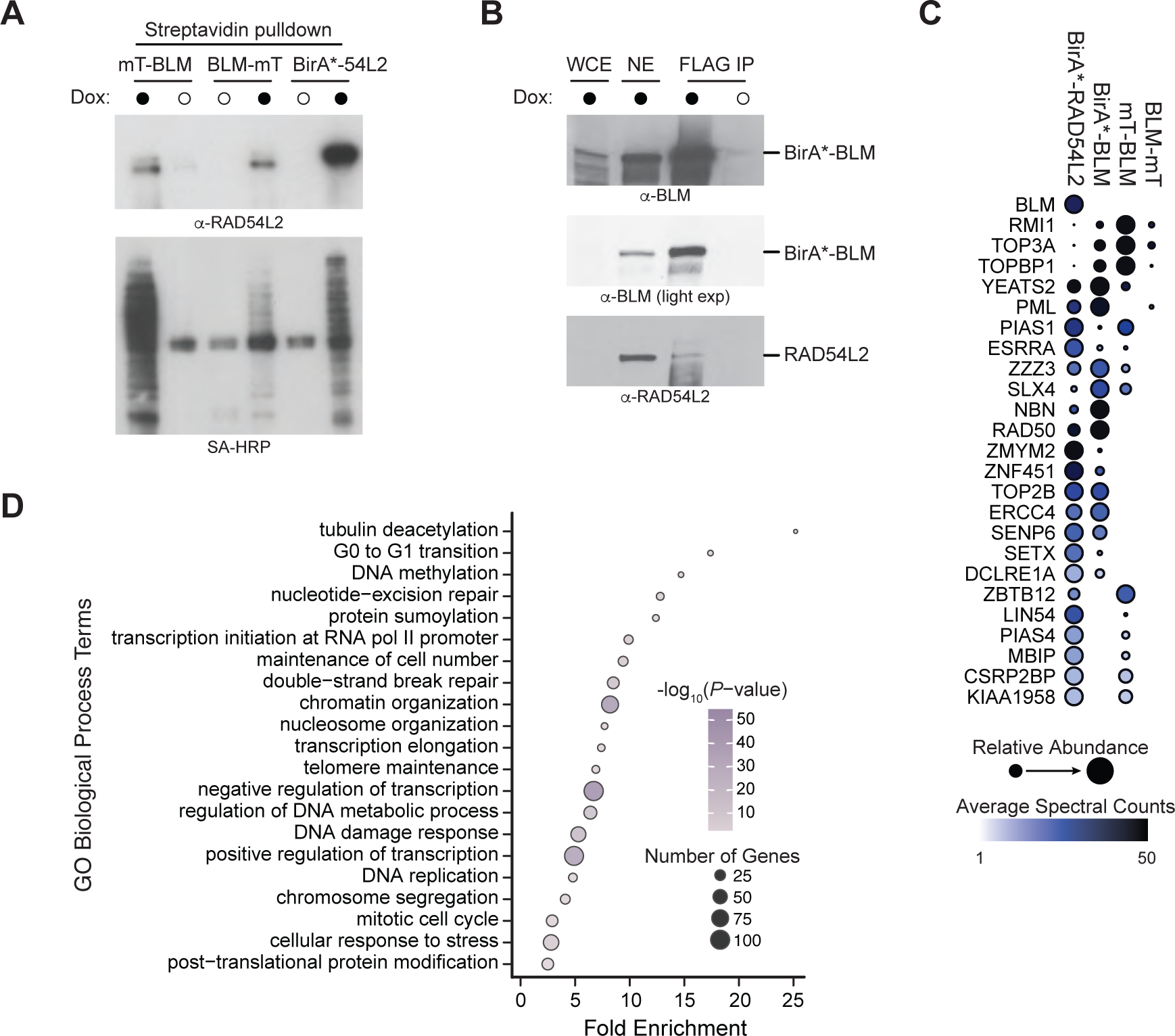
The RAD54L2 proximity interactome. A. Extracts of cells expressing mT-BLM, BLM-mT, or BirA*-RAD54L2, either without (open circles) or with (closed circles) doxycycline induction were affinity-purified with streptavidin-agarose. The affinity-purified biotinylated proteins were fractionated on SDS-PAGE and immunoblots were probed with anti-RAD54L2 antibodies or with streptavidin-HRP. B. Nuclear extracts (NE) of cells expressing mT-BLM, either without (open circles) or with (closed circles) doxycycline induction were subjected to affinity purification with anti-FLAG-agarose. The affinity-purified biotinylated proteins were fractionated on SDS-PAGE and immunoblots were probed with anti-FLAG (to detect mT-BLM), anti-TOP3A, or anti-RAD54L2 antibodies. The whole cell extract (WCE) is shown for comparison. C. Dot plot showing prey proteins identified with BirA*-RAD54L2, with the BLM BioID data from Fig. 2B plotted for comparison. High-confidence proximal proteins (BFDR ≤ 0.01) identified with RAD54L2 and at least one of the BLM fusions are shown. The RAD54L2 BioID screen was performed in quadruplicate, with dot colour indicating the average spectral counts. The dot size indicates relative abundance across the baits. D. Gene ontology (GO) biological process analysis for 238 high-confidence RAD54L2 proximal proteins identified in the BirA*-RAD54L2 BioID screen. The-fold enrichment for each GO term is indicated, the colours indicate the corrected P-values, and the size of the circles corresponds to the number of RAD54L2 proximal proteins annotated to the given GO term.

To gain further insight into RAD54L2 function we performed BioID with RAD54L2 (Fig 4C and Source Data 4C). Among the most prominent high-confidence proximal interactions was BLM. There was extensive overlap with the BLM proximity interactome, including TOP3A, RMI1, TOPBP1, SLX4, RAD50, and 19 other proteins (Fig 4C). Examining the functional enrichment of the high-confidence RAD54L2 proximal interactome (237 proteins), we found functions in common with the BLM interactome, including ‘DNA damage response’, ‘chromosome segregation’, and ‘protein sumoylation’ (Fig 4D, Source Data 2C, and Source Data 4D). We also noted functions consistent with roles in modulating chromatin accessibility, including ‘regulation of transcription’, ‘nucleosome organization’, and ‘transcription initiation’ (Fig 4D). We infer that RAD54L2 could perform distinct functions in chromatin structure and DNA repair, and that proximity to BLM could be a major aspect of RAD54L2 function.

### RAD54L2 promotes BLM focus formation

To develop mechanistic insight into how RAD54L2 promotes non-crossover recombination repair, we measured recruitment of BLM to sites of DNA replication stress, where BTRR function is important to re-start replication forks (Chan *et al*, 2007; Shorrocks *et al*, 2021). Association of BTRR with chromatin manifests as punctate BLM foci that are infrequent in unperturbed cells but are induced in the presence of the replication stressor hydroxyurea (Fig 5A and 5B). When *RAD54L2* was knocked out, BLM foci were absent, indicating that *RAD54L2* promotes the recruitment of BTRR to chromatin during DNA replication stress (Fig 5A and 5B). BLM functions at distinct stages of recombination repair of DSBs and stressed replication forks: BLM promotes long-range DNA resection to generate the template for RAD51-ssDNA filament formation (Gravel *et al*, 2008; Nimonkar *et al*, 2011) and it catalyzes dissolution of double Holliday junctions to suppress formation of crossover repair products (Wu & Hickson, 2003; Wu *et al*, 2006; Raynard *et al*, 2006; Bussen *et al*, 2007). We tested whether RAD54L2 was promoting BLM function during the early steps of recombination repair by measuring the recruitment of the resection protein MRE11 and the RAD51 recombinase to chromatin during DNA replication stress (Fig 5C and 5D). Cells that were deficient in RAD54L2 showed no changes in recruitment of resection or recombination proteins, indicating that RAD54L2 functions downstream of the RAD51 step of recombination repair.

**Figure 5.**
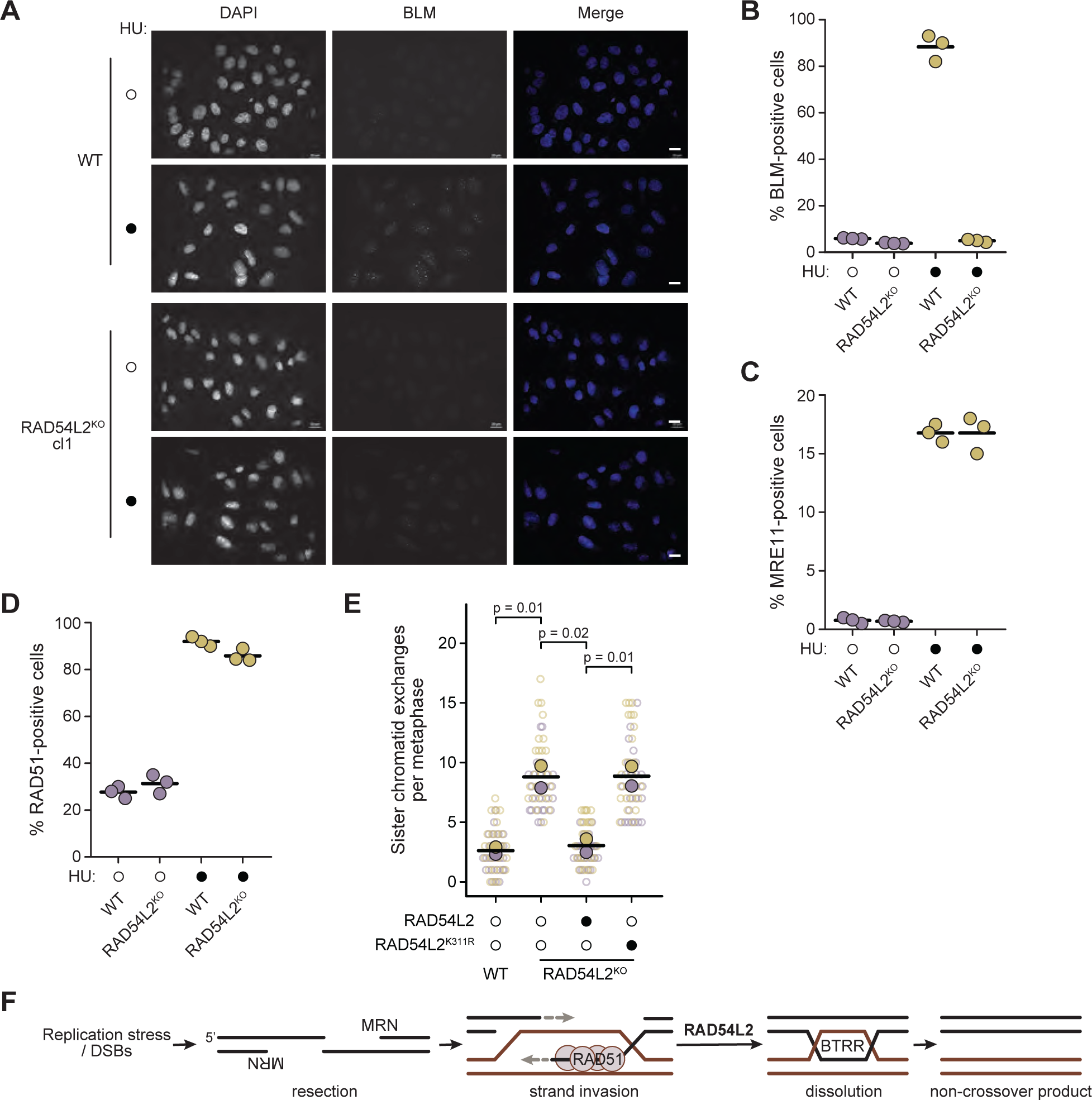
Recruitment of BLM, but not MRE11 or RAD51, to chromatin requires RAD54L2. A. Fluorescence micrographs of parental (WT) and RAD54L2 knockout cells, untreated (open circles) or treated with HU (closed circles). Images of cells stained with DAPI to illuminate the nuclear DNA, with antibodies to BLM, and the merged images are shown. Scale bars are 20 µm. B, C, D. The percent of parental (WT) and RAD54L2 knockout cells displaying BLM (B), MRE11 (C), or RAD51 (D) nuclear foci following control (open circles) or HU treatment (closed circles) is plotted for three replicates. Horizontal bars indicate the means. E. The number of SCEs per metaphase is plotted for two independent replicates for parental (WT) and RAD54L2 knockout cells. Where indicated by closed circles, cells were transfected with RAD54L2 or RAD54L2K311R The replicates are indicated by the different colours and the mean SCEs for each replicate is indicated by the filled circles. The p-values from one-sided Student’s t-tests comparing the replicate means are indicated. F. Model of the role of RAD54L2 in dissolution of DNA repair intermediates by BTRR. See text for details.

### RAD54L2 ATPase is required to suppress crossover recombination

SNF2-family proteins like RAD54L2 often utilize a conserved ATPase domain to modulate chromatin accessibility (Dürr *et al*, 2006). RAD54L2 ATPase activity requires intact Walker A and Walker B motifs (Walker *et al*, 1982), and ATPase activity is eliminated by changing the lysine residue in the Walker A motif to alanine (Rouleau *et al*, 2002). We tested if the integrity of the RAD54L2 ATPase domain was important for suppressing SCEs by transfecting *RAD54L2^KO^* cells with wild type or Walker A mutant (K311R) *RAD54L2*, and measuring SCEs (Fig 5E, EV5A, and EV5B). Expression of wild type *RAD54L2* reduced the amount of SCEs in *RAD54L2^KO^* cells to that seen in the parental U2OS cells. By contrast, the K311R mutant had no effect on SCEs in *RAD54L2^KO^*cells. RAD54L2 and RAD54L2^K311R^ proteins expressed well, although the K311R protein level was somewhat lower than the wild type (Fig EV5A and EV5B). We infer that RAD54L2 plays a catalytic role in suppressing SCEs rather than acting exclusively as a scaffold for BTRR recruitment to chromatin.

## Discussion

We present a comprehensive proximity interactome for the BLM-TOP3A-RMI1-RMI2 complex and highlight the functional importance of the SNF2-family ATPase RAD54L2 in promoting non-crossover recombination repair, likely by facilitating the association of BTRR with DNA damage sites on chromatin. Our data are consistent with a model (Fig 5F) whereby RAD54L2 promotes BTRR dissolution of late recombination intermediates resulting from double-strand DNA breaks or DNA replication fork re-start. RAD54L2 function appears to occur downstream of the MRE11 nuclease and the RAD51 recombinase and requires the ATPase activity of RAD54L2. The precise biochemical role of RAD54L2 in promoting non-crossover recombination outcomes catalyzed by BTRR awaits future investigation.

Given the distinct roles of the BTRR complex at different DNA structures, including DNA replication forks (Davies *et al*, 2007), double Holliday junctions (Wu *et al*, 2006; Raynard *et al*, 2006), ultra-fine anaphase bridges (Chan *et al*, 2007), D-loops (Bachrati *et al*, 2006; Harami *et al*, 2022), and G-quadruplexes (Sun *et al*, 1998), a comprehensive interactome for the BTRR complex offers a rich dataset for identifying proteins involved in the maintenance of genome stability. Our BioID studies complement existing affinity-purification/mass spectrometry datasets by capturing BTRR interactions in their native, unperturbed cellular environment. Given that affinity-based identification captures interactions that are stable to cell lysis conditions and are preserved or formed at the time of purification, there is usually only a modest overlap between AP-MS and BioID-MS datasets for DNA replication and repair proteins. For example, SLX4 affinity-purification and proximity-labeling interactomes had only 7.2% overlap (Aprosoff *et al*, 2023), and PCNA interactomes showed only 11.6% overlap (Srivastava *et al*, 2018). Our BLM proximity interactome, from three distinct BLM BioID experiments and comprising 101 high-confidence proximity interactions, shared 25 proteins with the 183 protein BLM interactome annotated in BioGRID (Oughtred *et al*, 2021) (14%). Likewise, we found that our RAD54L2 high-confidence (FDR≤0.01) proximity interactome of 237 proteins has little overlap with the 78 RAD54L2 protein interactors annotated in BioGRID (9 proteins, 11.5%) or with a recent RAD54L2 AP-MS analysis (3 proteins, 1.2%) (D’Alessandro *et al*, 2023). These data indicate the potential for proximity labeling methods to reveal interacting partners that are not detected using affinity purification methods.

It is, of course, important to evaluate whether a given interactome captures functional information. We find that the BTRR proximity interactome contains proteins that are known to function in concert with BTRR, including FANCM (Deans & West, 2009; Hoadley *et al*, 2012), additional BRAFT complex members FANCA and FANCG (Meetei *et al*, 2003), BRIP1/FANCJ (Suhasini *et al*, 2011), FANCD2 (Chaudhury *et al*, 2013), the RPA heterotrimer (RPA1, RPA2, and RPA3) (Yang *et al*, 2012; Xue *et al*, 2013; Doherty *et al*, 2005), TOPBP1 (Blackford *et al*, 2015), MLH1 (Pedrazzi *et al*, 2001), TP53 (Wang *et al*, 2001), BRCA1 (Acharya *et al*, 2014), RAD50 (Franchitto & Pichierri, 2002), and WRN (von Kobbe *et al*, 2002). As expected, the BTRR proximity interactome is strongly enriched for genes annotated to DNA replication, DNA recombination, and DNA repair gene ontology processes, and interestingly, functional enrichments are still apparent within the BTRR interactome even after removing the known BTRR interactors (Fig 1D). Additionally, we analysed the functions of 18 proteins in the BTRR interactome in suppressing sister chromatid exchanges and identified 15 that resulted in increased SCEs when depleted (Fig 3B). SCE suppressors include members of the FA complex (FANCM, FAAP24, FANCD2, CENPX), a replication protein (MCM10), ubiquitylation and sumoylation regulators (ABRAXAS2 (Feng *et al*, 2010; Zhang *et al*, 2014), UBR5, USP33, PIAS1), a histone reader (ZZZ3 (Mi *et al*, 2018)), and genome stability proteins (RAD54L2 (D’Alessandro *et al*, 2023; Zhang *et al*, 2023), SLF2 (Räschle *et al*, 2015), RADX (Dungrawala *et al*, 2017). The increases in SCEs that we found upon knockdown of FANCM, FAAP24, FANCD2, and SLF2 are similar to those previously reported following siRNA depletion of FANCM (Deans & West, 2009; Wang *et al*, 2013b), knockout of FAAP24 (Wang *et al*, 2013b), knockout of FANCD2 (Yamamoto *et al*, 2005), and in SLF2 patient fibroblasts (Grange *et al*, 2022). We conclude that the BTRR proximity interactome is rich in functional information.

Of the proteins in the BTRR proximity interactome, we found that RAD54L2 was biotinylated *in vivo* by both BirA*-BLM and miniTurbo-BLM (Fig 1B, 2B, and 4A). Both BLM fusion proteins could also immunoprecipitate RAD54L2 from nuclear or whole cell extracts (Fig 4B and Appendix Figure S1), suggesting that BLM and RAD54L2 form a stable complex. Knockdown or knockout of *RAD54L2* resulted in increased SCEs and decreased crossover recombination, indicating that *RAD54L2* deficiency phenocopies loss of *BLM*. Three lines of evidence suggest that *RAD54L2* is functioning in the *BLM* pathway to suppress SCEs. First, combining *BLM* knockdown with *RAD54L2* deficiency resulted in a small increase in SCEs that was neither additive nor multiplicative (Fig 3E), indicating that *BLM* and *RAD54L2* are in the same genetic pathway. Second, there is substantial overlap between the BLM and the RAD54L2 proximity interactomes (Fig 4C), consistent with BLM and RAD54L2 functioning in concert. Finally, cells deficient in *RAD54L2* fail to form BLM foci on chromatin in response to DNA replication stress (Fig 5A and 5B), suggesting that *RAD54L2* is important for recruitment of BLM to late recombination intermediates. We also found that the ATPase activity of RAD54L2 is important for its ability to suppress SCEs (Fig 5E), indicating a catalytic rather than a simple scaffolding function. Recent studies have implicated RAD54L2 in resistance to the topoisomerase poison etoposide (Zhang *et al*, 2023; D’Alessandro *et al*, 2023). RAD54L2 is proposed to function in concert with the SUMO E3 ligase ZNF451 to remove trapped TOP2 cleavage complexes from DNA. Interestingly, ZNF451 was among the high-confidence proximity interactions in our RAD54L2 BioID data (Fig 4C) and in our BLM and RMI1 BioID screens (Fig 1A). It remains to be seen whether ZNF451 participates in RAD54L2-mediated suppression of SCEs. RAD54L2 was associated with poor clinical outcomes in gastrointestinal stromal tumors (Schoppmann *et al*, 2013) and in a pediatric AML cohort treated with etoposide (Nguyen *et al*, 2023). Given that RAD54L2 is a potential prognostic biomarker and drug target, the RAD54L2 and BLM proximity interactomes could have clinical utility.

## Materials and Methods

### Plasmids

To construct plasmids for generating stable BirA* cell lines for BioID, *BLM* and *RMI2* cDNAs were amplified from pCR4-TOPO-BLM and pOTB7-RMI2 (MGC Project Team *et al*, 2009). The PCR products were digested with AscI and XhoI and cloned into pcDNA5/FRT/TO-Flag-BirA* (Lambert *et al*, 2015) to yield pcDNA5/FRT/TO-FLAG-BirA*-BLM and pcDNA5/FRT/TO-FLAG-BirA*-RMI2. pcDNA5/FRT/TO-Flag-TOP3A and pcDNA5/FRT/TO-Flag-RMI1 (Yang *et al*, 2012) were digested with AscI and XhoI and cloned into pcDNA5/FRT/TO-Flag-BirA* to yield pcDNA5/FRT/TO-FLAG-BirA*-TOP3A and pcDNA5/FRT/TO-FLAG-BirA*-RMI1. *RAD54L2* cDNA from Mammalian Gene Collection (MGC Project Team *et al*, 2009) was cloned into pcDNA5/FRT/TO-Flag-BirA* to yield pcDNA5/FRT/TO-FLAG-BirA*-RAD54L2.

Plasmids for miniTurbo (Branon *et al*, 2018) were constructed with BLM from pcDNA3/BLM (Gaymes *et al*, 2002), cloned into pDONR201 to yield pDONR201-BLM. The BLM cDNA was moved from pDONR201-BLM into pDEST-miniTurbo-Nterm and pDEST-miniTurbo-Cterm (Branon *et al*, 2018) obtained from the Gingras Lab, using a Gateway LR reaction to yield pDEST-miniTurbo-Nterm-BLM and pDEST-miniTurbo-Cterm-BLM.

For rescue experiments, the *RAD54L2* cDNA was assembled from a partial cDNA clone encoding Met109 to Lys1467 was from the human ORFeome collection (Lamesch *et al*, 2007) and a synthetic DNA fragment encoding Met1 to Glu108 (Invitrogen). *RAD54L2^K311R^* was constructed by QuikChange site-directed mutagenesis with primers (5’-AGATCACTTGCAAAGTTCTCCCCAGACCCATGCTG-3’ and 5’-CAGCATGGGTCTGGGGAGAACTTTGCAAGTGATCT-3’). Both constructs were confirmed by whole plasmid sequencing.

To generate RAD54L2 knockouts a double-stranded DNA oligo encoding a sgRNA targeting the RAD54L2 5’ region from the TKOv3 library (Hart *et al*, 2017) (5’-AAGATGGGCAGCAGCCGCCG CGG–3’) was cloned into pSpCas9(BB)-2A-Puro (PX459) (RRID: Addgene_48139) to yield PX459-RAD54L2.

All constructs were confirmed by Sanger sequencing unless otherwise noted.

### Human cell culture

Flp-In-T-REx-293 (RRID: CVCL_U427) cells were grown in Dulbecco’s modified Eagle’s medium (DMEM; Wisent #319-005-CL) supplemented with 10% fetal bovine serum (FBS; Wisent #098-150), 1% penicillin/streptomycin (Wisent #450-201-EL), 15 μg/mL blasticidin (BioShop #BLA477.25) and 100 μg/mL zeocin (InvivoGen #ant-zn-05). Stably transfected Flp-In-T-REx-293 lines were cultured with 15 μg/mL blasticidin (BioShop #BLA477.25) and 200 μg/mL hygromycin (BioShop # HYG002.202). U2OS (RRID: CVCL_0042) and U2OS DR-GFP (RRID:CVCL_B0A7) cells were grown in McCoy’s 5A medium (Gibco # 16600082) supplemented with 10% fetal bovine serum (FBS; Wisent #098-150) and 1% penicillin/streptomycin (Wisent #450-201-EL). All cell lines were grown at 37°C and 5% CO_2_. Cell lines were routinely monitored to ensure the absence of *Mycoplasma* contamination using the MycoAlert PLUS Mycoplasma Detection Kit (Lonza #LT07-705).

### Production of stable cell lines for BioID and miniTurbo proximity-dependent biotinylation

To generate stable cell lines, Flp-In-T-REx-293 cells were transfected with the BioID or miniTurbo construct of interest plus the Flp recombinase expression vector pOG44, using Lipofectamine 3000 (ThermoFisher # L3000001) or X-tremeGENE HP (Roche # 6366236001), according to the manufacturer’s instructions. At 24h post-transfection, cells were expanded for selection with 200 μg/mL hygromycin B. Hygromycin-resistant cells were expanded in the presence of 200 μg/mL hygromycin to create polyclonal stable cell lines. Where indicated, expression of fusion proteins was induced for 24h by the addition of doxycycline (Sigma-Aldrich # D9891) to 5 µg/ml. The expression of BirA* and miniTurbo fusion proteins was confirmed by immunoblotting (Expanded View Figures EV1 and EV2).

### Streptavidin purification of biotinylated proteins and on-bead trypsin digest

Proximal proteins of BLM, TOP3A, RMI1 and RMI2 were identified as previously described (Lambert *et al*, 2015). Proximal proteins in RAD54L2 BioID and BLM miniTurbo were prepared as previously described (St-Germain *et al*, 2020). For each biological replicate, HEK293 cells stably expressing a tetracycline regulated BirA* (two 150 mm dishes) or miniTurbo fusion protein (five 150 mm dishes) were grown to 60-80% confluency. Cells were treated with 1 μg/mL doxycycline and 50 μM biotin (Sigma-Aldrich # B-4639) for 24 hours for BirA* fusions or 30 minutes for miniTurbo fusions. Medium was removed and cells were washed twice with cold PBS. Cells were scraped and pooled together and washed two times with cold PBS. Cell pellets were flash frozen and stored at −80°C.

The BirA* expressing cells from two 150 mm dishes were lysed at 1:10 cell pellet weight (g): RIPA buffer volume (mL) (50 mM Tris-HCl pH 7.5, 150 mM NaCl, 1% Triton X-100, 1 mM EDTA, 1 mM EGTA, 0.1% SDS, Sigma protease inhibitors P8340 1:500, and 0.5% sodium deoxycholate), for 1 hour on a nutator at 4°C. For miniTurbo fusions, five 150 mm dishes were incubated at 4°C in 10 mL RIPA buffer, for 1 hour on a nutator. After incubation, 1µl of benzonase (250U) was added to each sample and the lysates were sonicated (3 x 10 second bursts with 2 seconds rest in between) on ice at 65% amplitude. The lysates were then centrifuged for 30 min at 20,817 x g at 4 °C. During this step, streptavidin-sepharose beads (GE # 17-5113-01) were washed 3 times with 1 mL RIPA buffer (minus protease inhibitors and sodium deoxycholate). Beads were pelleted at 400 x g for 1 minute in between washes. After centrifugation, supernatants from the clarified lysates were transferred to 15 mL falcon tubes and a 30 µL bed volume of washed beads was added to each. Affinity purification was performed at 4°C on a nutator for 3 hours, then beads were pelleted (400 x g, 1 min), the supernatant removed, and the beads transferred to a 1.5 mL Eppendorf tube in 1 mL RIPA buffer (minus protease inhibitors and sodium deoxycholate). The beads were washed by pipetting up and down (four times per wash step) with 1 mL RIPA buffer (minus protease inhibitors and sodium deoxycholate), followed by two washes in TAP lysis buffer (50 mM HEPES-KOH pH 8.0, 100 mM KCl, 10% glycerol, 2 mM EDTA, 0.1% NP-40), then 3 washes in 50 mM ammonium bicarbonate pH 8 (ABC). Beads were pelleted by centrifugation (400 x g, 1 min) and the supernatant aspirated between wash steps. After the last wash, all residual 50 mM ABC was pipetted off.

Beads were re-suspended in 30 µL (for BioID) and 200 µL (for miniTurbo) of 50 mM ammonium bicarbonate (ABC) pH 8 with 1µg trypsin added and incubated at 37°C overnight with mixing on a rotating disc. The next day, an additional 0.5 µg of trypsin was added to each sample (in 10 µL 50 mM ABC) and the samples incubated for an additional 2 hours at 37°C with mixing on a nutator. Beads were pelleted (400 x g, 2 min) and the supernatant transferred to a fresh 1.5 mL Eppendorf tube. The beads were then rinsed 2 times with 30 µL of HPLC H_2_O for BirA* fusions or 150 µl of 50 mM ABC for miniTurbo fusions each time (pelleting beads at 400 x g, 2 min in between) and these rinses combined with the original supernatant. The pooled supernatant was acidified by adding 50% formic acid (FA) to a final concentration of 2% v/v and dried by vacuum centrifugation. For BirA* fusions, dried samples were resuspended in 12 µL of 5% FA.

### Mass spectrometric analysis

Affinity-purified digested material in 12 µL of 5% formic acid was centrifuged at 16,100 x g for 1 minute before 6 µL was loaded onto silica columns pre-packed with 10-12 cm of C18 reversed-phase material (ZorbaxSB, 3.5 µm). The loaded column was placed in-line with an LTQ-Orbitrap Velos or an Orbitrap Elite (Thermo Electron, Bremen, Germany) equipped with a nanoelectrospray ion source (Proxeon Biosystems, Odense, Denmark) connected in-line to a NanoLC-Ultra 2D plus HPLC system (Eksigent, Dublin, USA). The LTQ-Orbitrap Velos or Orbitrap Elite instrument under Xcalibur 2.0 was operated in the data-dependent mode to automatically switch between MS and up to 10 subsequent MS/MS acquisitions. Buffer A is 100 parts H_2_O, 0.1 part formic acid; buffer B is 100 parts acetonitrile, 0.1 part formic acid. The HPLC gradient program delivered an acetonitrile gradient over 125 minutes. For the first twenty minutes, the flow rate was 400 µL/min at 2%B. The flow rate was then reduced to 200 µL/min, and the fraction of solvent B increased in a linear fashion to 35% at 95.5 minutes. Solvent B was then increased to 80% over 5 minutes and maintained at that level until 107 minutes. The mobile phase was then reduced to 2% B until the end of the run (125 min). The parameters for data-dependent acquisition on the mass spectrometer were 1 centroid MS (mass range 400-2000) followed by MS/MS on the 10 most abundant ions. General parameters were: activation type = CID, isolation width = 1 m/z, normalized collision energy = 35, activation Q = 0.25, activation time = 10 msec. For data-dependent acquisition, the minimum threshold was 500, the repeat count = 1, the repeat duration = 30 sec, the exclusion size list = 500, exclusion duration = 30 sec, exclusion mass width (by mass) = low 0.03, high 0.03.

### Mass spectrometry data analysis

All mass spectrometry data files were stored, searched, and analyzed using ProHits 7.0 Laboratory Information Management System (LIMS) platform (Liu *et al*, 2016). Within ProHits, RAW files were converted to an MGF format using ProteoWizard (V3.0.10702). The data were searched using Mascot V2.3.02 (Perkins *et al*, 1999) and Comet V2018.01 rev.4 (Eng *et al*, 2013). The spectra were searched with the human and adenovirus sequences in the NCBI Reference Sequence Database (version 57, January 30, 2013), supplemented with “common contaminants” from the Max Planck Institute (http://maxquant.org/contaminants.zip) and the Global Proteome Machine (GPM; ftp://ftp.thegpm.org/fasta/cRAP/crap.fasta), forward and reverse sequences (labeled “gi|9999” or “DECOY”), sequence tags, and streptavidin, for a total of 72,482 entries. Database parameters were set to search for tryptic cleavages, allowing up to 2 missed cleavages per peptide with a mass tolerance of 12 ppm for precursors with charges of 2+ to 4+ and a tolerance of 0.6Da for fragment ions. Variable modifications were selected for deamidated asparagine and glutamine, and oxidated methionine. Results from each search engine were analyzed through TPP (the Trans-Proteomic Pipeline, v.4.7 POLAR VORTEX rev 1) via the iProphet pipeline (Shteynberg *et al*, 2011). All proteins with an iProphet probability ≥95% were used for analysis.

### Protein proximity interaction scoring

Significance analysis of interactome express (SAINTexpress) version 3.6.3 was used to calculate the probability of potential associations/interactions compared to background contaminants using default parameters (Teo *et al*, 2014). In brief, SAINTexpress is a statistical tool that compares the spectral counts of each prey identified with a given biotin ligase-tagged bait against a set of negative controls. For BioID, negative controls consisted of streptavidin affinity purifications from cells expressing BirA*-FLAG and BirA*, 6 biological replicates each. Two or more biological replicates were collected for all cell lines and conditions. For BioID proximity interaction scoring, two replicates with the highest spectral counts for each prey were used for baits; four replicates were used for negative controls for SAINTexpress. SAINT scores were averaged across replicates, and these averages were used to calculate a Bayesian false discovery rate (BFDR); preys with BFDR ≤1% were considered high-confidence protein interactions. All non-human protein interactors (did not start with “NP” in Prey column) were removed from the SAINT analysis. Proximity interaction scores are tabulated in SourceData1AB.xlsx, SourceData2B.xlsx, and SourceData4C.xlsx.

### Gene Ontology term enrichment analysis

GO enrichment analysis was performed using high-confidence proximity interactors as inputs for the Princeton Generic Gene Ontology (GO) Term Finder (Boyle *et al*, 2004). Redundant GO terms were consolidated with REVIGO (Supek *et al*, 2011) using a medium list size and the SimRel semantic similarity measure, and in some cases manually, and terms that included more than 10% of genes were removed. The complete unfiltered outputs from GOTermFinder are provided in SourceData1D.xlsx, SourceData2C.xlsx, and SourceData4D.xlsx.

### Protein-protein interaction data mining

Physical interactions with low-throughput or high-throughput evidence for BTRR complex members were downloaded from BioGRID (https://thebiogrid.org/; accessed 24/09/2023; SourceData1AB.xlsx). The physical interaction network for BTRR BioID hits (Fig 1C) was generated with GeneMANIA (https://genemania.org/; accessed 23/01/2024; SourceData1C.xlsx). The BLM proximity interaction network was generated from the miniTurbo data using Cytoscape 3.10.0 (Shannon *et al*, 2003).

### Gene knockdowns with RNA interference

For the assays in Fig 3, U2OS cells were transfected with small interfering RNA (siRNA) pools (Dharmacon/Horizon) or individual siRNAs, as listed in the Reagents and Tools Table. An siRNA that does not have a target in the human genome was used as the negative control. All experiments were performed 48 h after siRNA transfection to achieve optimal protein depletion, unless indicated otherwise. Lipofectamine RNAiMAX (Invitrogen) was used to carry out siRNA transfections.

### Sister chromatid exchange assays

Approximately 0.2 x 10^6^ U2OS cells per well were seeded and grown overnight. U2OS cells were then cultured for 48 hours (approximately two rounds of DNA replication) in medium supplemented with 10 μM bromodeoxyuridine (BrdU; Sigma-Aldrich # B5002) to differentially label sister chromatids. Cells were treated with 100 ng/µl colcemid (Gibco # 15212012) for 2 hours to arrest the cells in metaphase. The cells were harvested, resuspended in 10 mL 0.075M KCl, and incubated at 37°C for 20 minutes. The swollen cells were pelleted by centrifugation, the supernatant was removed, and the cells were resuspended 10 mL of fresh 3:1 methanol:acetic acid fixative solution. The cells were washed with the fixative solution twice. After the last fixative wash, the cells were stored in 1 mL of the fixative solution. Microscope slides were cleaned with ethanol, dried, and steamed for a few seconds above a beaker of boiling water immediately before chromosome spreading. The cell suspension was dropped onto the steamed slide until the surface was covered, slides were air dried, and quickly dipped in 70% acetic acid. Slides were then cured in the dark for 24 hours at room temperature. Cells were stained with 0.75 μg/ml Hoechst 3342 for 30 min and exposed to UV light for 30 min, washed with 2X SSC buffer for 15 min, and stained with 0.1 µg/ml DAPI in PBS. The resulting metaphase chromosome spreads were imaged on a Zeiss AxioImager Z1 widefield fluorescence microscope equipped with DAPI filter cube 49. A minimum of 25 mitoses were analyzed per experiment. Statistical support was evaluated with the Student’s t-test.

### Direct-repeat GFP recombination assays

U2OS cells carrying the DR-GFP reporter (Pierce *et al*, 1999) were transfected with siRNAs as indicated above. After 24 h, cells were transfected with the I-SceI expression plasmid pCBASceI(Richardson *et al*, 1998) using X-tremeGENE HP. 48 h after I-SceI transfection, cells were harvested and resuspended in PBS buffer (Ca^2+^/ Mg^2+^ Free) plus 0.5% heat-inactivated FBS. Live cells were harvested in FACS buffer (Ca^2+^/ Mg^2+^-free PBS with 0.5% heat-inactivated FBS) and incubated on ice during data acquisition. Flow cytometry to identify GFP-positive cells was performed on BD LSR II Flow Cytometer, and 10 000 events were acquired based on the first FSC/SSC gate. Flow cytometry data were analyzed using FlowJo (BD Biosciences, RRID: SCR_008520). For experiments with RAD54L2 knockout lines, cells were transfected with PX459-RAD54L2 and PX459-K311R and grown for 24h before proceeding with pCBASceI transfection. Statistical support was evaluated with the Student’s t-test.

### Construction of RAD54L2 knockout U2OS lines by CRISPR-CAS editing

U2OS or U2OS DR-GFP cells were transfected with PX459-sgRAD54L2 using X-tremeGENE HP and selected with 1.5 μg/ml puromycin for 72 hours. Individual cells were isolated by flow cytometry, expanded, and screened for RAD54L2 expression by western blotting (Expanded View Fig EV4A and EV4C). Editing of the RAD54L2 genomic locus was confirmed by amplifying approximately 250bp upstream and downstream of the Cas9 cut site from genomic DNA followed by Sanger sequencing. The Cas9 edits in RAD54L2 alleles were inferred from the Sanger sequencing data using ICE(Conant *et al*, 2022):

**Summary Table:** Cas9 edits in RAD54L2 alleles.

| Summary Table: Cas9 edits in RAD54L2 alleles |  |  |
| --- | --- | --- |
| RAD54L2 KO | KO Score | Indels |
| U2OS cl.1 | 98 | {'-14': 98.0} |
| U2OS cl.2 | 71 | {'7': 2.0, '-1': 52.0, '-4': 3.0, '-7': 4.0, '-8': 1.0, '10': 2.0, '12': 1.0, '19': 3.0, '-11': 1.0, '-17': 1.0, '-28': 2.0} |
| DR-GFP cl.3 | 97 | {'1': 65.0, '-1': 32.0} |
| DR-GFP cl.1 | 96 | {'1': 31.0, '2': 33.0, '-10': 32.0} |

### Antibodies

Primary antibodies used in this study were: mouse anti-FLAG M2 (Sigma-Aldrich #F3165, RRID: AB_259529, 1:1000-1:2500), mouse anti-*α*-tubulin (Cell Signaling Technologies #3873, RRID: AB_1904178, 1:5000), mouse anti-GAPDH (Cell Signaling Technologies #97166, RRID: AB_2756824, 1:5000), mouse anti-V5 (Cell Signaling Technologies #D3H8Q; 1:1000), anti-BLM (Cell Signaling Technologies #2742; Bethyl Laboratories #A300-120A; 1:1000 for immunofluorescence), anti-ABRAXAS2 (Abcam #ab68801; 1:1000), anti-RAD54L2 (ABclonal #A6144, 1:1000), anti-ZZZ3 (Bethyl Laboratories #A303-331A; 1:1000), anti-TOP3A (Proteintech # 14525-1-AP; 1:1000), anti-RMI1 (Raised in house #6534, 1:500), anti-RAD51 (Abcam #ab133534), anti-MRE11(Novus Biologicals #NB100-473, 1:1000). Secondary antibodies used for immunoblotting were: goat anti-mouse IgG HRP (Invitrogen # 31430, 1:5000) and goat anti-rabbit IgG HRP (Invitrogen # 31460, 1:5000). Secondary antibody used for immunofluorescence were Donkey anti-Goat IgG, Alexa Fluor 546 (Invitrogen # A-11056, 1:1000) and Donkey anti-Rabbit IgG, Alexa Fluor 488 (Invitrogen # A-21206, 1:1000), and Donkey anti-Mouse IgG, Alexa Fluor 488 (Invitrogen # A-21202, 1:1000).

Streptavidin-HRP (Cell Signaling Technologies #3999; 1:5000) was used to detect biotinylated proteins.

### Immunoblotting and immunoprecipitation

Whole-cell extracts for immunoblotting were prepared by lysing 1 - 2 × 10^6^ cells in RIPA buffer (50 mM Tris pH 7.5, 150 mM NaCl, 0.1% SDS, 2 mM EDTA, 0.5% sodium deoxycholate, 1% (v/v) Triton X-100) containing 1x cOmplete EDTA-free Protease Inhibitor Cocktail (Roche) and 1x PhosSTOP phosphatase inhibitor (Roche) for 30 minutes at 4°C with gentle agitation. Extracts were clarified by centrifugation at 14 000 rpm for 15 minutes at 4°C, and the supernatant was aliquoted into new 1.5 mL tubes. Protein concentrations were measured using the Pierce Rapid Gold BCA Protein Assay Kit (ThermoFisher #A53225). Whole cell extracts (20-50 *μ*g) were diluted in 4x Laemmli sample buffer (250 mM Tris pH 6.8, 5% SDS, 40% (w/v) glycerol, 0.02% bromophenol blue, 10% *β*-mercaptoethanol) and boiled at 95°C for 5 minutes prior to fractionation by SDS-PAGE. Proteins were transferred to nitrocellulose membranes, blocked for one hour in 5% (w/v) skim milk in TBS with 0.2% Tween-20 (TBST) and probed with primary antibodies overnight at 4°C. Membranes were washed three times in TBST for five minutes each, probed with secondary antibodies for one hour, then washed another three times in TBST for five minutes each. Proteins were detected using SuperSignal West Pico PLUS chemiluminescent substrate (ThermoFisher #34580) and visualized on X-ray film.

Nuclear extracts for co-immunoprecipitations of BirA*-BLM and RAD54L2 were prepared by resuspending cell pellets in cytoplasmic lysis buffer (50 mM Tris pH7.5, 10 mM NaCl, 1.5 mM MgCl_2_, 10% glycerol, 0.34 M sucrose, 0.1% Triton X-100), harvesting nuclei by centrifugation at 14 000 rpm for 10 minutes at 4°C, followed by resuspending in nuclear extraction buffer (50 mM Tris pH7.5, 250 mM NaCl, 10% glycerol, 1 mM EDTA). The nuclear extract was clarified by centrifugation at 14 000 rpm for 10 minutes at 4°, and the supernatant (nuclear extract) was transferred to a fresh tube.

For co-immunoprecipitations, whole cell or nuclear extracts were incubated with 2.5 µg of anti-FLAG antibody bound to 50 µl Dynabeads Protein G (Invitrogen #10003D) overnight at 4°C with gentle rotation. Immunoprecipitates were washed three times with cytoplasmic lysis buffer and eluted in 2X Laemmli buffer (4% SDS, 120mM Tris-HCl, 0.005% bromophenol blue, 20% glycerol, 200 mM DTT) at 95°C for 10 min. S. Proteins were fractionated by SDS-PAGE and analyzed by immunoblotting as described above.

### Immunofluorescence microscopy

For the detection of BLM, MRE11, and RAD51 foci, 0.4×10^6^ cells parental and *RAD54L2*-deficient U2OS cells were seeded were seeded onto HCl acid-etched coverslips and cultured in the presence or absence of 4 mM HU (BioBasic) for 24h before fixation. Cells were washed with ice-cold PBS, and pre-extracted with CSK buffer (300 mM sucrose, 100 mM NaCl, 3 mM MgCl_2_, 10 mM PIPES pH 7.0, 0.5% (v/v) Triton X-100) for 15 minutes on ice. Extracted cells were fixed with 4% (w/v) PFA in PBS for 10 minutes at room temperature, washed three times in PBS, and incubated in blocking buffer (10% goat serum (ThermoFisher #16210064), 0.5% NP-40, 5% (w/v) saponin in PBS) 30 min. Blocking buffer containing the appropriate primary antibodies was added to the cover slips for 2 hours, the slips were washed three times in PBS, and incubated in blocking buffer containing secondary antibodies and 0.5 *μ*g/mL 4’,6-diamidino-2-phenylindole (DAPI; Sigma-Aldrich #D9542) for one hour, protected from light. After washing with PBS, coverslips were mounted with ProLong Gold Antifade (ThermoFisher #P36930). Images were acquired on a Zeiss AxioObserver Z1 confocal microscope with a 63x oil-immersion objective lens. Cells that were positive for BLM, MRE11, or RAD51 foci were quantified by manual counting. At least 50 cells per condition were analyzed in each experiment.

## Data Availability

Data has been deposited as a complete submission to the MassIVE repository (https://massive.ucsd.edu/ProteoSAFe/static/massive.jsp) and assigned the accession number MSV000093876. The ProteomeXchange accession is PXD048612. The dataset is currently available for reviewers at ftp://MSV000093876@massive.ucsd.edu. Please login with username MSV000093876_reviewer; password: BLMhelicase. The dataset will be made public upon acceptance of the manuscript. Raw and processed source data for all figures is included in the Source Data files.

## Supporting information

Reagents and Tools Table

## Acknowledgements

We thank Dan Durocher for providing cell lines. ACG is a Tier 1 Canada Research Chair in Functional Proteomics. Proteomics work was performed at the Network Biology Collaborative Centre at the Lunenfeld-Tanenbaum Research Institute, a facility supported by Canada Foundation for Innovation funding, by the Ontarian Government, by Genome Canada, and Ontario Genomics (OGI-139). GWB is a Tier I Canada Research Chair in Genome Integrity and was supported by the Canadian Institutes of Health Research (FDN-159913). We are grateful to work on the lands of the Mississaugas of the Credit, the Anishnaabeg, the Haudenosaunee, and the Wendat peoples, land that is now home to many diverse First Nations, Inuit, and Métis peoples.

## Author Contributions

JJH: Formal analysis, Investigation, Methodology, Formal analysis, Writing-original draft, Writing-review & editing. EC: Investigation, Writing-review & editing. WHD: Investigation. CJW: Investigation, Formal analysis, Writing-review & editing. JS-G: Investigation, Formal analysis, Writing-review & editing. BR: Formal analysis, Funding acquisition, Supervision. ACG: Formal analysis, Funding acquisition, Supervision, Writing-review & editing. GWB: Conceptualization, Formal analysis, Funding acquisition, Supervision, Writing-original draft, Writing-review & editing.

## Expanded View Figure Legends

**Figure EV1.**
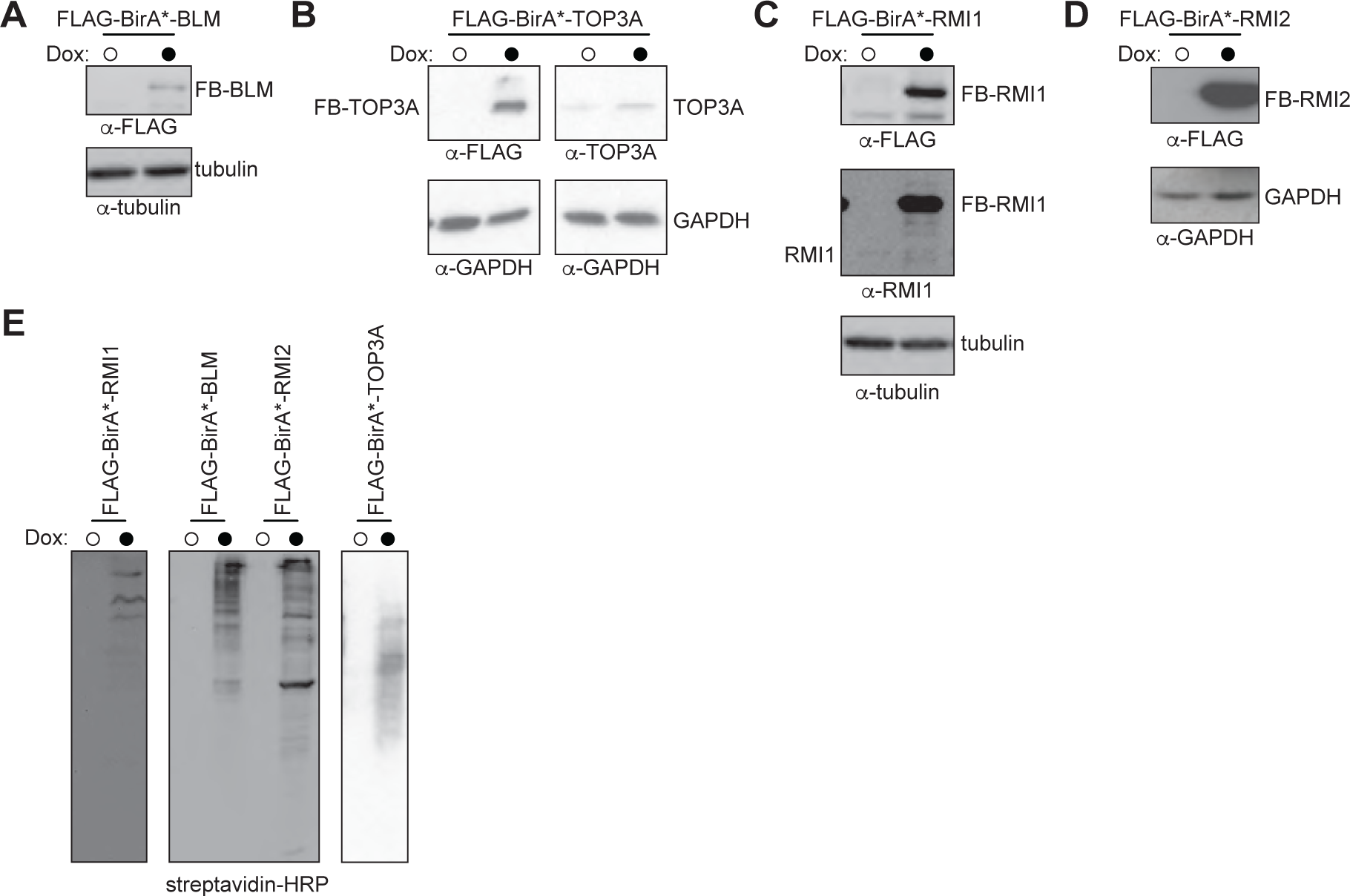
Expression of active BTRR BirA* fusion proteins. A. The stable FLAG-BirA*-BLM Flp-In T-Rex HEK293 cell line was treated with doxycycline (closed circle) or vehicle (open circle) to induce expression of FLAG-BirA*-BLM (FB-BLM) and subjected to immunoblot analysis, probing with anti-FLAG or anti-tubulin antibodies, as indicated. B. The stable FLAG-BirA*-TOP3A Flp-In T-Rex HEK293 cell line was treated with doxycycline (closed circle) or vehicle (open circle) to induce expression of FLAG-BirA*-TOP3A (FB-TOP3A) and subjected to immunoblot analysis, probing with anti-FLAG, anti-TOP3A, or anti-GAPDH antibodies, as indicated. C. The stable FLAG-BirA*-RMI1 Flp-In T-Rex HEK293 cell line was treated with doxycycline (closed circle) or vehicle (open circle) to induce expression of FLAG-BirA*-RMI1 (FB-RMI1) and subjected to immunoblot analysis, probing with anti-FLAG, anti-RMI1, or anti-tubulin antibodies, as indicated. D. The stable FLAG-BirA*-RMI2 Flp-In T-Rex HEK293 cell line was treated with doxycycline (closed circle) or vehicle (open circle) to induce expression of FLAG-BirA*-RMI2 (FB-RMI2) and subjected to immunoblot analysis, probing with anti-FLAG, or anti-GAPDH antibodies, as indicated. E. Stable Flp-In T-Rex HEK293 cell lines were treated with doxycycline (closed circle) or vehicle (open circle) to induce expression of the indicated FLAG-BirA* fusion proteins. Extracts of the cells were fractionated by SDS-PAGE, and biotinylated proteins were detected with streptavidin-HRP.

**Figure EV2.**
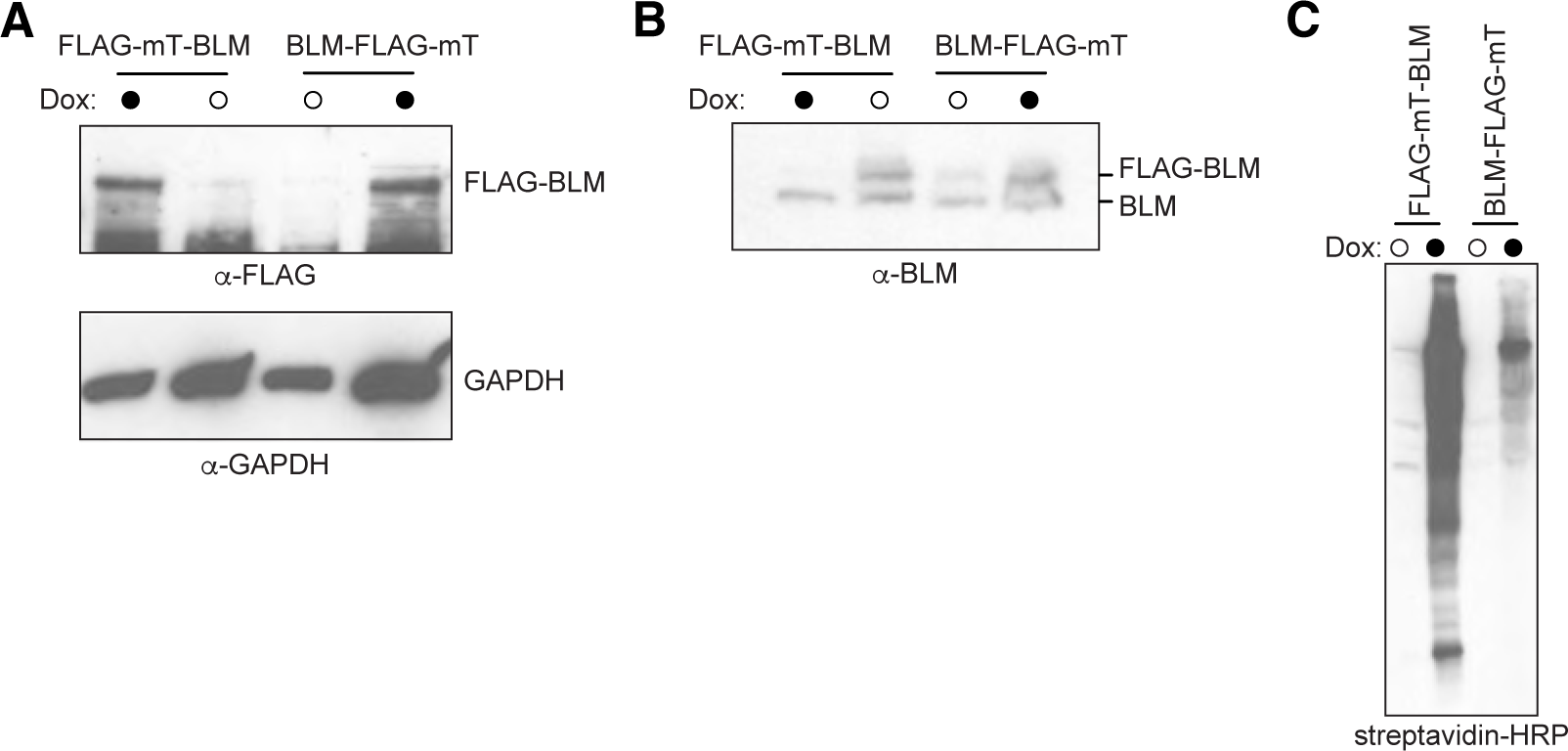
Expression of active BLM miniTurbo fusions. A. The stable FLAG-miniTurbo-BLM and BLM-FLAG-miniTurbo cell lines were treated with doxycycline (closed circle) or vehicle (open circle) to induce expression of FLAG-BirA*-BLM (FB-BLM) and subjected to immunoblot analysis, probing with anti-FLAG or anti-GAPDH antibodies, as indicated. B. The stable FLAG-miniTurbo-BLM and BLM-FLAG-miniTurbo cell lines were treated with doxycycline (closed circle) or vehicle (open circle) to induce expression of FLAG-BirA*-BLM (FB-BLM) and subjected to immunoblot analysis, probing with anti-BLM antibodies. C. The stable FLAG-miniTurbo-BLM and BLM-FLAG-miniTurbo cell lines were treated with doxycycline (closed circle) or vehicle (open circle) to induce expression of FLAG-BirA*-BLM. Extracts of the cells were fractionated by SDS-PAGE, and biotinylated proteins were detected with streptavidin-HRP.

**Figure EV3.**
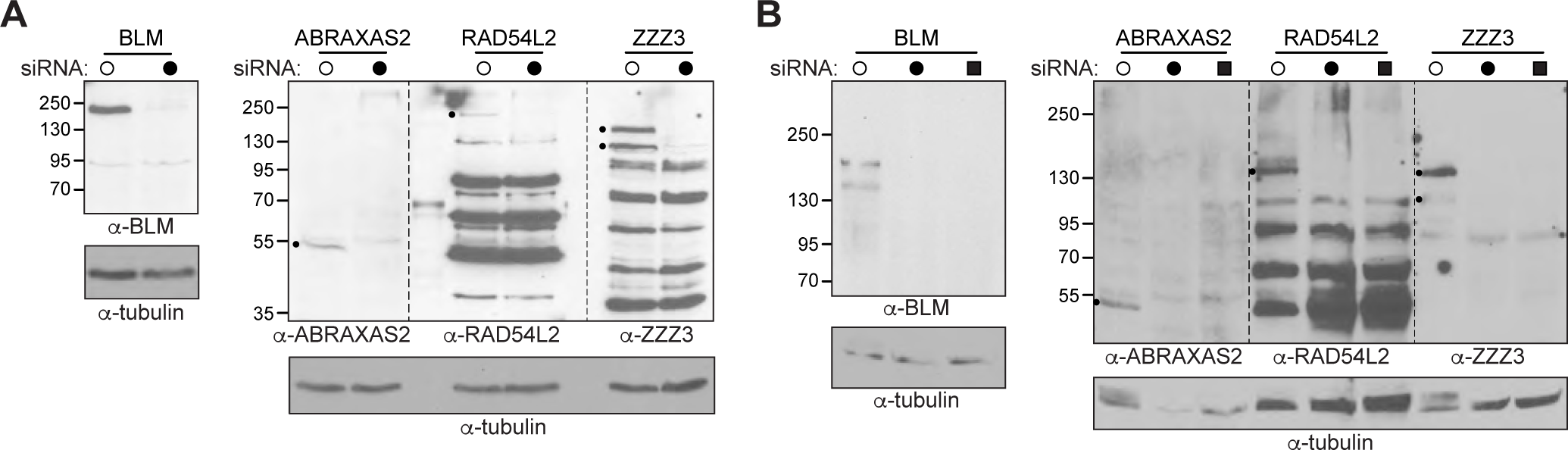
Knockdown of BLM, ABRAXAS2, RAD54L2, and ZZZ3. A. Control (open circles) or the indicated siRNAs (closed circles) were transfected into U2OS cells. After 48h, protein depletion was examined by immunoblot analysis, probing with the indicated antibodies. Small circles mark putative RAD54L2 and ZZZ3 polypeptides. B. Control (open circles), the indicated siRNAs (closed circles), or the indicated siRNAs and the I-SceI expression plasmid (closed squares) were transfected into U2OS DR-GFP cells. After 48h, protein depletion was examined by immunoblot analysis, probing with the indicated antibodies. Small circles mark putative RAD54L2 and ZZZ3 polypeptides.

**Figure EV4.**
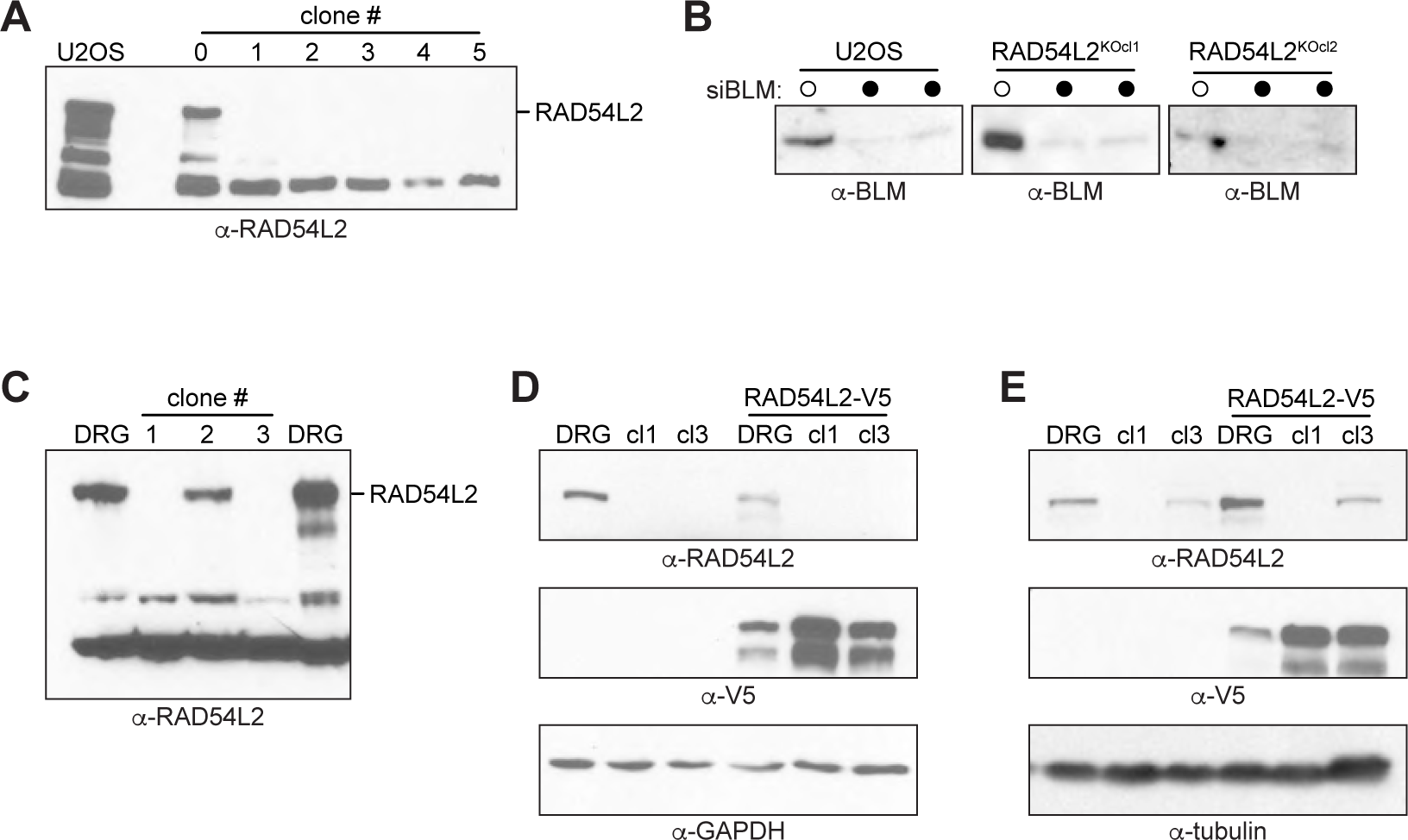
CRISPR/Cas disruption of *RAD54L2.* A. U2OS cells were transfected with a plasmid expressing Cas9 and an sgRNA targeting *RAD54L2*, cloned, and examined by immunoblot analysis, probing with antibodies against RAD54L2. Parental cells (U2OS) were also examined. The position of RAD54L2 is indicated. B. U2OS cells and two *RAD54L2* knockout lines (cl1, cl2) were treated with control (open circles) or *BLM* siRNAs (closed circles), and examined by immunoblot analysis, probing with antibodies against BLM. Two replicates are shown for each BLM knockdown. C. U2OS DR-GFP cells were transfected with a plasmid expressing Cas9 and an sgRNA targeting *RAD54L2*, cloned, and examined by immunoblot analysis, probing with antibodies against RAD54L2. Parental cells (DRG) were also examined. The position of RAD54L2 is indicated. D. U2OS DR-GFP (DRG) and two U2OS DR-GFP *RAD54L2* knockout lines (cl1, cl3) were examined by immunoblotting, probing with the indicated antibodies. Where indicated, the cells were transfected with a plasmid carrying *RAD54L2* tagged with the V5 epitope (RAD54L2-V5). E. As in panel D, for the independent experimental replicate.

**Figure EV5.**
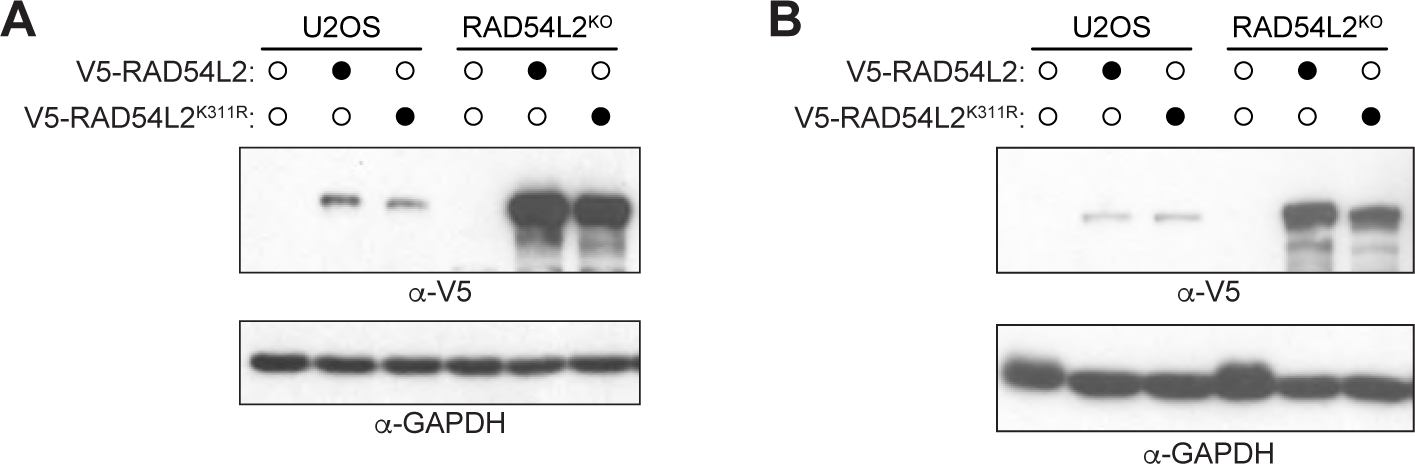
Rescue of *RAD54L2* knockouts. A. U2OS cells or RAD54L2-deficient U2OS cells (clone 1) were mock-transfected (open circles) or transfected with V5-tagged RAD54L2 or RAD54L2K311R (closed circles). Extracts of the cells were examined by immunoblot analysis, probing with antibodies against V5 or GAPDH. B. As in panel A, for the independent experimental replicate.

## Appendix Figure Legends

**Figure S1.**
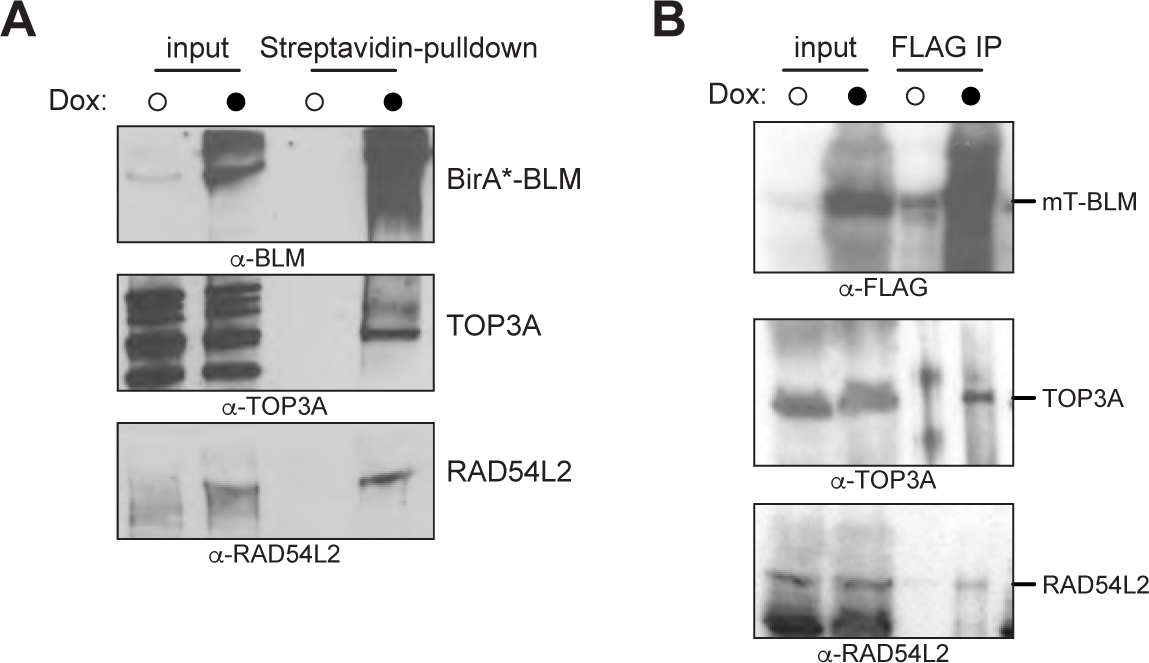
Co-affinity-purification of BLM and RAD54L2. A. Extracts of cells expressing BirA*-BLM, either without (open circles) or with (closed circles) doxycycline induction, were affinity-purified with streptavidin-agarose. The affinity-purified biotinylated proteins were fractionated on SDS-PAGE and immunoblots were probed with anti-BLM, anti-TOP3A, or anti-RAD54L2 antibodies. B. Extracts of cells expressing mT-BLM, either without (open circles) or with (closed circles) doxycycline induction were subjected to affinity purification with anti-FLAG-agarose to purify mT-BLM. The affinity-purified proteins were fractionated on SDS-PAGE and immunoblots were probed with anti-FLAG, anti-TOP3A, or anti-RAD54L2 antibodies.

## References

Ababou M (2021) Bloom syndrome and the underlying causes of genetic instability. Molecular Genetics and Metabolism 133: 35–48

Acharya S, Kaul Z, Gocha AS, Martinez AR, Harris J, Parvin JD & Groden J (2014) Association of BLM and BRCA1 during Telomere Maintenance in ALT Cells. PLOS ONE 9: e103819

Aprosoff CM, Dyakov BJA, Cheung VHW, Wong CJ, Palandra M, Gingras A-C & Wyatt HDM (2023) Comprehensive Interactome Mapping of the DNA Repair Scaffold SLX4 Using Proximity Labeling and Affinity Purification. J Proteome Res 22: 1660–1681

Bachrati CZ, Borts RH & Hickson ID (2006) Mobile D-loops are a preferred substrate for the Bloom’s syndrome helicase. Nucleic Acids Res 34: 2269–2279

Bass TE, Luzwick JW, Kavanaugh G, Carroll C, Dungrawala H, Glick GG, Feldkamp MD, Putney R, Chazin WJ & Cortez D (2016) ETAA1 acts at stalled replication forks to maintain genome integrity. Nat Cell Biol 18: 1185–1195

Belhadj S, Khurram A, Bandlamudi C, Palou-Márquez G, Ravichandran V, Steinsnyder Z, Wildman T, Catchings A, Kemel Y, Mukherjee S, et al (2023) NBN Pathogenic Germline Variants are Associated with Pan-Cancer Susceptibility and In Vitro DNA Damage Response Defects. Clinical Cancer Research 29: 422–431

Bhattacharyya S, Keirsey J, Russell B, Kavecansky J, Lillard-Wetherell K, Tahmaseb K, Turchi JJ & Groden J (2009) Telomerase-associated Protein 1, HSP90, and Topoisomerase IIα Associate Directly with the BLM Helicase in Immortalized Cells Using ALT and Modulate Its Helicase Activity Using Telomeric DNA Substrates *. Journal of Biological Chemistry 284: 14966–14977

Bizard AH & Hickson ID (2014) The dissolution of double Holliday junctions. Cold Spring Harb Perspect Biol 6: a016477

Blackford AN, Nieminuszczy J, Schwab RA, Galanty Y, Jackson SP & Niedzwiedz W (2015) TopBP1 interacts with BLM to maintain genome stability but is dispensable for preventing BLM degradation. Mol Cell 57: 1133–1141

Boyle EI, Weng S, Gollub J, Jin H, Botstein D, Cherry JM & Sherlock G (2004) GO::TermFinder--open source software for accessing Gene Ontology information and finding significantly enriched Gene Ontology terms associated with a list of genes. Bioinformatics 20: 3710–3715

Branon TC, Bosch JA, Sanchez AD, Udeshi ND, Svinkina T, Carr SA, Feldman JL, Perrimon N & Ting AY (2018) Efficient proximity labeling in living cells and organisms with TurboID. Nat Biotechnol 36: 880–887

Bussen W, Raynard S, Busygina V, Singh AK & Sung P (2007) Holliday junction processing activity of the BLM-Topo IIIalpha-BLAP75 complex. J Biol Chem 282: 31484–31492

Bythell-Douglas R & Deans AJ (2021) A Structural Guide to the Bloom Syndrome Complex. Structure 29: 99–113

Chan K-L, North PS & Hickson ID (2007) BLM is required for faithful chromosome segregation and its localization defines a class of ultrafine anaphase bridges. EMBO J 26: 3397–3409

Chaudhury I, Sareen A, Raghunandan M & Sobeck A (2013) FANCD2 regulates BLM complex functions independently of FANCI to promote replication fork recovery. Nucleic Acids Res 41: 6444–6459

Cho NH, Cheveralls KC, Brunner A-D, Kim K, Michaelis AC, Raghavan P, Kobayashi H, Savy L, Li JY, Canaj H, et al (2022) OpenCell: Endogenous tagging for the cartography of human cellular organization. Science 375: eabi6983

Conant D, Hsiau T, Rossi N, Oki J, Maures T, Waite K, Yang J, Joshi S, Kelso R, Holden K, et al (2022) Inference of CRISPR Edits from Sanger Trace Data. The CRISPR Journal 5: 123–130

Conrad S, Künzel J & Löbrich M (2011) Sister chromatid exchanges occur in G2-irradiated cells. Cell Cycle 10: 222–228

Coulthard R, Deans AJ, Swuec P, Bowles M, Costa A, West SC & McDonald NQ (2013) Architecture and DNA Recognition Elements of the Fanconi Anemia FANCM-FAAP24 Complex. Structure 21: 1648–1658

D’Alessandro G, Morales-Juarez DA, Richards SL, Nitiss KC, Serrano-Benitez A, Wang J, Thomas JC, Gupta V, Voigt A, Belotserkovskaya R, et al (2023) RAD54L2 counters TOP2-DNA adducts to promote genome stability. Science Advances 9: eadl2108

Danino YM, Molitor L, Rosenbaum-Cohen T, Kaiser S, Cohen Y, Porat Z, Marmor-Kollet H, Katina C, Savidor A, Rotkopf R, et al (2023) BLM helicase protein negatively regulates stress granule formation through unwinding RNA G-quadruplex structures. Nucleic Acids Research: gkad613

Davalos AR, Kaminker P, Hansen RK & Campisi J (2004) ATR and ATM-dependent movement of BLM helicase during replication stress ensures optimal ATM activation and 53BP1 focus formation. Cell Cycle 3: 1579–1586

Davies SL, North PS, Dart A, Lakin ND & Hickson ID (2004) Phosphorylation of the Bloom’s syndrome helicase and its role in recovery from S-phase arrest. Mol Cell Biol 24: 1279–1291

Davies SL, North PS & Hickson ID (2007) Role for BLM in replication-fork restart and suppression of origin firing after replicative stress. Nat Struct Mol Biol 14: 677–679

Deans AJ & West SC (2009) FANCM connects the genome instability disorders Bloom’s Syndrome and Fanconi Anemia. Mol Cell 36: 943–953

Doherty KM, Sommers JA, Gray MD, Lee JW, von Kobbe C, Thoma NH, Kureekattil RP, Kenny MK & Brosh RM (2005) Physical and functional mapping of the replication protein a interaction domain of the werner and bloom syndrome helicases. J Biol Chem 280: 29494–29505

Drosopoulos WC, Kosiyatrakul ST & Schildkraut CL (2015) BLM helicase facilitates telomere replication during leading strand synthesis of telomeres. J Cell Biol 210: 191–208

Dungrawala H, Bhat KP, Le Meur R, Chazin WJ, Ding X, Sharan SK, Wessel SR, Sathe AA, Zhao R & Cortez D (2017) RADX Promotes Genome Stability and Modulates Chemosensitivity by Regulating RAD51 at Replication Forks. Mol Cell 67: 374–386.e5

Dürr H, Flaus A, Owen-Hughes T & Hopfner K-P (2006) Snf2 family ATPases and DExx box helicases: differences and unifying concepts from high-resolution crystal structures. Nucleic Acids Res 34: 4160–4167

Eladad S, Ye T-Z, Hu P, Leversha M, Beresten S, Matunis MJ & Ellis NA (2005) Intra-nuclear trafficking of the BLM helicase to DNA damage-induced foci is regulated by SUMO modification. Human Molecular Genetics 14: 1351–1365

Eng JK, Jahan TA & Hoopmann MR (2013) Comet: an open-source MS/MS sequence database search tool. Proteomics 13: 22–24

Feng L, Wang J & Chen J (2010) The Lys63-specific deubiquitinating enzyme BRCC36 is regulated by two scaffold proteins localizing in different subcellular compartments. J Biol Chem 285: 30982–30988

Franchitto A & Pichierri P (2002) Bloom’s syndrome protein is required for correct relocalization of RAD50/MRE11/NBS1 complex after replication fork arrest. J Cell Biol 157: 19–30

Franz M, Rodriguez H, Lopes C, Zuberi K, Montojo J, Bader GD & Morris Q (2018) GeneMANIA update 2018. Nucleic Acids Res 46: W60–W64

Gaymes TJ, North PS, Brady N, Hickson ID, Mufti GJ & Rassool FV (2002) Increased error-prone non homologous DNA end-joining--a proposed mechanism of chromosomal instability in Bloom’s syndrome. Oncogene 21: 2525–2533

German J (1997) Bloom’s syndrome. XX. The first 100 cancers. Cancer Genetics and Cytogenetics 93: 100–106

German J, Schonberg S, Louie E & Chaganti RS (1977) Bloom’s syndrome. IV. Sister-chromatid exchanges in lymphocytes. Am J Hum Genet 29: 248–255

Gönenc II, Elcioglu NH, Martinez Grijalva C, Aras S, Großmann N, Praulich I, Altmüller J, Kaulfuß S, Li Y, Nürnberg P, et al (2022) Phenotypic spectrum of BLM-and RMI1-related Bloom syndrome. Clinical Genetics 101: 559–564

Grange LJ, Reynolds JJ, Ullah F, Isidor B, Shearer RF, Latypova X, Baxley RM, Oliver AW, Ganesh A, Cooke SL, et al (2022) Pathogenic variants in SLF2 and SMC5 cause segmented chromosomes and mosaic variegated hyperploidy. Nat Commun 13: 6664

Gravel S, Chapman JR, Magill C & Jackson SP (2008) DNA helicases Sgs1 and BLM promote DNA double-strand break resection. Genes Dev 22: 2767–2772

Guo Y, Xu H, Huang M & Ruan Y (2023) BLM promotes malignancy in PCa by inducing KRAS expression and RhoA suppression via its interaction with HDGF and activation of MAPK/ERK pathway. J Cell Commun Signal 17: 757–772

Harami GM, Pálinkás J, Seol Y, Kovács ZJ, Gyimesi M, Harami-Papp H, Neuman KC & Kovács M (2022) The toposiomerase IIIalpha-RMI1-RMI2 complex orients human Bloom’s syndrome helicase for efficient disruption of D-loops. Nat Commun 13: 654

Hart T, Tong AHY, Chan K, Van Leeuwen J, Seetharaman A, Aregger M, Chandrashekhar M, Hustedt N, Seth S, Noonan A, et al (2017) Evaluation and Design of Genome-Wide CRISPR/SpCas9 Knockout Screens. G3 (Bethesda) 7: 2719–2727

Heijink AM, Stok C, Porubsky D, Manolika EM, de Kanter JK, Kok YP, Everts M, de Boer HR, Audrey A, Bakker FJ, et al (2022) Sister chromatid exchanges induced by perturbed replication can form independently of BRCA1, BRCA2 and RAD51. Nat Commun 13: 6722

Hein MY, Hubner NC, Poser I, Cox J, Nagaraj N, Toyoda Y, Gak IA, Weisswange I, Mansfeld J, Buchholz F, et al (2015) A human interactome in three quantitative dimensions organized by stoichiometries and abundances. Cell 163: 712–723

Hoadley KA, Xu D, Xue Y, Satyshur KA, Wang W & Keck JL (2010) Structure and cellular roles of the RMI core complex from the Bloom syndrome dissolvasome. Structure 18: 1149–1158

Hoadley KA, Xue Y, Ling C, Takata M, Wang W & Keck JL (2012) Defining the molecular interface that connects the Fanconi anemia protein FANCM to the Bloom syndrome dissolvasome. Proc Natl Acad Sci U S A 109: 4437–4442

Hu P, Beresten SF, van Brabant AJ, Ye TZ, Pandolfi PP, Johnson FB, Guarente L & Ellis NA (2001) Evidence for BLM and Topoisomerase IIIalpha interaction in genomic stability. Hum Mol Genet 10: 1287–1298

Hudson DF, Amor DJ, Boys A, Butler K, Williams L, Zhang T & Kalitsis P (2016) Loss of RMI2 Increases Genome Instability and Causes a Bloom-Like Syndrome. PLOS Genetics 12: e1006483

von Kobbe C, Karmakar P, Dawut L, Opresko P, Zeng X, Brosh RM, Hickson ID & Bohr VA (2002) Colocalization, physical, and functional interaction between Werner and Bloom syndrome proteins. J Biol Chem 277: 22035–22044

Lambert J-P, Tucholska M, Go C, Knight JDR & Gingras A-C (2015) Proximity biotinylation and affinity purification are complementary approaches for the interactome mapping of chromatin-associated protein complexes. J Proteomics 118: 81–94

Lamesch P, Li N, Milstein S, Fan C, Hao T, Szabo G, Hu Z, Venkatesan K, Bethel G, Martin P, et al (2007) hORFeome v3.1: a resource of human open reading frames representing over 10,000 human genes. Genomics 89: 307–315

Langlois RG, Bigbee WL, Jensen RH & German J (1989) Evidence for increased in vivo mutation and somatic recombination in Bloom’s syndrome. Proc Natl Acad Sci U S A 86: 670–674

LaRocque JR, Stark JM, Oh J, Bojilova E, Yusa K, Horie K, Takeda J & Jasin M (2011) Interhomolog recombination and loss of heterozygosity in wild-type and Bloom syndrome helicase (BLM)-deficient mammalian cells. Proc Natl Acad Sci U S A 108: 11971–11976

Liu G, Knight JDR, Zhang JP, Tsou C-C, Wang J, Lambert J-P, Larsen B, Tyers M, Raught B, Bandeira N, et al (2016) Data Independent Acquisition analysis in ProHits 4.0. J Proteomics 149: 64–68

Lönn U, Lönn S, Nylen U, Winblad G & German J (1990) An abnormal profile of DNA replication intermediates in Bloom’s syndrome. Cancer Res 50: 3141–3145

Lu R, O’Rourke JJ, Sobinoff AP, Allen JAM, Nelson CB, Tomlinson CG, Lee M, Reddel RR, Deans AJ & Pickett HA (2019) The FANCM-BLM-TOP3A-RMI complex suppresses alternative lengthening of telomeres (ALT). Nat Commun 10: 2252

Martin C-A, Sarlós K, Logan CV, Thakur RS, Parry DA, Bizard AH, Leitch A, Cleal L, Ali NS, Al-Owain MA, et al (2018) Mutations in TOP3A Cause a Bloom Syndrome-like Disorder. Am J Hum Genet 103: 221–231

Meetei AR, Sechi S, Wallisch M, Yang D, Young MK, Joenje H, Hoatlin ME & Wang W (2003) A Multiprotein Nuclear Complex Connects Fanconi Anemia and Bloom Syndrome. Molecular and Cellular Biology 23: 3417–3426

MGC Project Team, Temple G, Gerhard DS, Rasooly R, Feingold EA, Good PJ, Robinson C, Mandich A, Derge JG, Lewis J, et al (2009) The completion of the Mammalian Gene Collection (MGC). Genome Res 19: 2324–2333

Mi W, Zhang Y, Lyu J, Wang X, Tong Q, Peng D, Xue Y, Tencer AH, Wen H, Li W, et al (2018) The ZZ-type zinc finger of ZZZ3 modulates the ATAC complex-mediated histone acetylation and gene activation. Nat Commun 9: 3759

Nguyen NHK, Rafiee R, Tagmount A, Sobh A, Loguinov A, de Jesus Sosa AK, Elsayed AH, Gbadamosi M, Seligson N, Cogle CR, et al (2023) Genome-wide CRISPR/Cas9 screen identifies etoposide response modulators associated with clinical outcomes in pediatric AML. Blood Advances 7: 1769–1783

Nimonkar AV, Genschel J, Kinoshita E, Polaczek P, Campbell JL, Wyman C, Modrich P & Kowalczykowski SC (2011) BLM-DNA2-RPA-MRN and EXO1-BLM-RPA-MRN constitute two DNA end resection machineries for human DNA break repair. Genes Dev 25: 350–362

Oughtred R, Rust J, Chang C, Breitkreutz B, Stark C, Willems A, Boucher L, Leung G, Kolas N, Zhang F, et al (2021) The BioGRID database: A comprehensive biomedical resource of curated protein, genetic, and chemical interactions. Protein Sci 30: 187–200

Pedrazzi G, Perrera C, Blaser H, Kuster P, Marra G, Davies SL, Ryu GH, Freire R, Hickson ID, Jiricny J, et al (2001) Direct association of Bloom’s syndrome gene product with the human mismatch repair protein MLH1. Nucleic Acids Res 29: 4378–4386

Perkins DN, Pappin DJ, Creasy DM & Cottrell JS (1999) Probability-based protein identification by searching sequence databases using mass spectrometry data. Electrophoresis 20: 3551–3567

Pierce AJ, Johnson RD, Thompson LH & Jasin M (1999) XRCC3 promotes homology-directed repair of DNA damage in mammalian cells. Genes Dev 13: 2633–2638

Räschle M, Smeenk G, Hansen RK, Temu T, Oka Y, Hein MY, Nagaraj N, Long DT, Walter JC, Hofmann K, et al (2015) DNA repair. Proteomics reveals dynamic assembly of repair complexes during bypass of DNA cross-links. Science 348: 1253671

Raynard S, Bussen W & Sung P (2006) A double Holliday junction dissolvasome comprising BLM, topoisomerase IIIalpha, and BLAP75. J Biol Chem 281: 13861–13864

de Renty C & Ellis NA (2017) Bloom’s Syndrome: Why Not Premature Aging? A comparison of the BLM and WRN helicases. Ageing Res Rev 33: 36–51

Renwick A, Thompson D, Seal S, Kelly P, Chagtai T, Ahmed M, North B, Jayatilake H, Barfoot R, Spanova K, et al (2006) ATM mutations that cause ataxia-telangiectasia are breast cancer susceptibility alleles. Nat Genet 38: 873–875

Richardson C, Moynahan ME & Jasin M (1998) Double-strand break repair by interchromosomal recombination: suppression of chromosomal translocations. Genes Dev 12: 3831–3842

Rouleau N, Domans’kyi A, Reeben M, Moilanen A-M, Havas K, Kang Z, Owen-Hughes T, Palvimo JJ & Jänne OA (2002) Novel ATPase of SNF2-like Protein Family Interacts with Androgen Receptor and Modulates Androgen-dependent Transcription. MBoC 13: 2106–2119

Roux KJ, Kim DI, Raida M & Burke B (2012) A promiscuous biotin ligase fusion protein identifies proximal and interacting proteins in mammalian cells. J Cell Biol 196: 801–810

Schoppmann SF, Vinatzer U, Popitsch N, Mittlböck M, Liebmann-Reindl S, Jomrich G, Streubel B & Birner P (2013) Novel clinically relevant genes in gastrointestinal stromal tumors identified by exome sequencing. Clin Cancer Res 19: 5329–5339

Sengupta S, Linke SP, Pedeux R, Yang Q, Farnsworth J, Garfield SH, Valerie K, Shay JW, Ellis NA, Wasylyk B, et al (2003) BLM helicase-dependent transport of p53 to sites of stalled DNA replication forks modulates homologous recombination. The EMBO Journal 22: 1210–1222

Shannon P, Markiel A, Ozier O, Baliga NS, Wang JT, Ramage D, Amin N, Schwikowski B & Ideker T (2003) Cytoscape: a software environment for integrated models of biomolecular interaction networks. Genome Res 13: 2498–2504

Shorrocks A-MK, Jones SE, Tsukada K, Morrow CA, Belblidia Z, Shen J, Vendrell I, Fischer R, Kessler BM & Blackford AN (2021) The Bloom syndrome complex senses RPA-coated single-stranded DNA to restart stalled replication forks. Nat Commun 12: 585

Shteynberg D, Deutsch EW, Lam H, Eng JK, Sun Z, Tasman N, Mendoza L, Moritz RL, Aebersold R & Nesvizhskii AI (2011) iProphet: multi-level integrative analysis of shotgun proteomic data improves peptide and protein identification rates and error estimates. Mol Cell Proteomics 10: M111.007690

Singh TR, Ali AM, Busygina V, Raynard S, Fan Q, Du C, Andreassen PR, Sung P & Meetei AR (2008) BLAP18/RMI2, a novel OB-fold-containing protein, is an essential component of the Bloom helicase-double Holliday junction dissolvasome. Genes Dev 22: 2856–2868

Srivastava M, Chen Z, Zhang H, Tang M, Wang C, Jung SY & Chen J (2018) Replisome Dynamics and Their Functional Relevance upon DNA Damage through the PCNA Interactome. Cell Reports 25: 3869–3883.e4

St-Germain JR, Samavarchi Tehrani P, Wong C, Larsen B, Gingras A-C & Raught B (2020) Variability in Streptavidin–Sepharose Matrix Quality Can Significantly Affect Proximity-Dependent Biotinylation (BioID) Data. J Proteome Res 19: 3554–3561

Sturzenegger A, Burdova K, Kanagaraj R, Levikova M, Pinto C, Cejka P & Janscak P (2014) DNA2 Cooperates with the WRN and BLM RecQ Helicases to Mediate Long-range DNA End Resection in Human Cells *. Journal of Biological Chemistry 289: 27314– 27326

Suhasini AN, Rawtani NA, Wu Y, Sommers JA, Sharma S, Mosedale G, North PS, Cantor SB, Hickson ID & Brosh RM (2011) Interaction between the helicases genetically linked to Fanconi anemia group J and Bloom’s syndrome. EMBO J 30: 692–705

Sun H, Karow JK, Hickson ID & Maizels N (1998) The Bloom’s syndrome helicase unwinds G4 DNA. J Biol Chem 273: 27587–27592

Supek F, Bošnjak M, Škunca N & Šmuc T (2011) REVIGO summarizes and visualizes long lists of gene ontology terms. PLoS One 6: e21800

Takata H, Nishijima H, Ogura S, Sakaguchi T, Bubulya PA, Mochizuki T & Shibahara K (2009) Proteome Analysis of Human Nuclear Insoluble Fractions. Genes Cells 14: 975–990

Teo G, Liu G, Zhang J, Nesvizhskii AI, Gingras A-C & Choi H (2014) SAINTexpress: improvements and additional features in Significance Analysis of Interactome software. J Proteomics 100: 37–43

Walker JE, Saraste M, Runswick MJ & Gay NJ (1982) Distantly related sequences in the alpha-and beta-subunits of ATP synthase, myosin, kinases and other ATP-requiring enzymes and a common nucleotide binding fold. EMBO J 1: 945–951

Wan L, Han J, Liu T, Dong S, Xie F, Chen H & Huang J (2013) Scaffolding protein SPIDR/KIAA0146 connects the Bloom syndrome helicase with homologous recombination repair. Proceedings of the National Academy of Sciences 110: 10646–10651

Wang J, Chen J & Gong Z (2013a) TopBP1 Controls BLM Protein Level to Maintain Genome Stability. Mol Cell 52: 10.1016/j.molcel.2013.10.012

Wang T, Hu J, Li Y, Bi L, Guo L, Jia X, Zhang X, Li D, Hou X-M, Modesti M, et al (2022) Bloom Syndrome Helicase Compresses Single-Stranded DNA into Phase-Separated Condensates. Angew Chem Int Ed Engl 61: e202209463

Wang XW, Tseng A, Ellis NA, Spillare EA, Linke SP, Robles AI, Seker H, Yang Q, Hu P, Beresten S, et al (2001) Functional interaction of p53 and BLM DNA helicase in apoptosis. J Biol Chem 276: 32948–32955

Wang Y, Leung JW, Jiang Y, Lowery MG, Do H, Vasquez KM, Chen J, Wang W & Li L (2013b) FANCM and FAAP24 maintain genome stability via cooperative as well as unique functions. Mol Cell 49: 997–1009

Weinstock DM, Nakanishi K, Helgadottir HR & Jasin M (2006) Assaying double-strand break repair pathway choice in mammalian cells using a targeted endonuclease or the RAG recombinase. Methods Enzymol 409: 524–540

van Wietmarschen N, Merzouk S, Halsema N, Spierings DCJ, Guryev V & Lansdorp PM (2018) BLM helicase suppresses recombination at G-quadruplex motifs in transcribed genes. Nat Commun 9: 271

Wu L, Bachrati CZ, Ou J, Xu C, Yin J, Chang M, Wang W, Li L, Brown GW & Hickson ID (2006) BLAP75/RMI1 promotes the BLM-dependent dissolution of homologous recombination intermediates. Proc Natl Acad Sci U S A 103: 4068–4073

Wu L, Davies SL, North PS, Goulaouic H, Riou JF, Turley H, Gatter KC & Hickson ID (2000) The Bloom’s syndrome gene product interacts with topoisomerase III. J Biol Chem 275: 9636–9644

Wu L & Hickson ID (2003) The Bloom’s syndrome helicase suppresses crossing over during homologous recombination. Nature 426: 870–874

Xue X, Raynard S, Busygina V, Singh AK & Sung P (2013) Role of replication protein A in double holliday junction dissolution mediated by the BLM-Topo IIIα-RMI1-RMI2 protein complex. J Biol Chem 288: 14221–14227

Yamamoto K, Hirano S, Ishiai M, Morishima K, Kitao H, Namikoshi K, Kimura M, Matsushita N, Arakawa H, Buerstedde J-M, et al (2005) Fanconi anemia protein FANCD2 promotes immunoglobulin gene conversion and DNA repair through a mechanism related to homologous recombination. Mol Cell Biol 25: 34–43

Yang J, O’Donnell L, Durocher D & Brown GW (2012) RMI1 promotes DNA replication fork progression and recovery from replication fork stress. Mol Cell Biol 32: 3054–3064

Yankiwski V, Noonan JP & Neff NF (2001) The C-terminal domain of the Bloom syndrome DNA helicase is essential for genomic stability. BMC Cell Biology 2: 11

Yin J, Sobeck A, Xu C, Meetei AR, Hoatlin M, Li L & Wang W (2005) BLAP75, an essential component of Bloom’s syndrome protein complexes that maintain genome integrity. The EMBO Journal 24: 1465–1476

Yusa K, Horie K, Kondoh G, Kouno M, Maeda Y, Kinoshita T & Takeda J (2004) Genome-wide phenotype analysis in ES cells by regulated disruption of Bloom’s syndrome gene. Nature 429: 896–899

Zhang H, Xiong Y, Sun Y, Park J-M, Su D, Feng X, Keast S, Tang M, Huang M, Wang C, et al (2023) RAD54L2-mediated DNA damage avoidance pathway specifically preserves genome integrity in response to topoisomerase 2 poisons. Sci Adv 9: eadi6681

Zhang J, Cao M, Dong J, Li C, Xu W, Zhan Y, Wang X, Yu M, Ge C, Ge Z, et al (2014) ABRO1 suppresses tumourigenesis and regulates the DNA damage response by stabilizing p53. Nat Commun 5: 5059

Zhao W, Vaithiyalingam S, San Filippo J, Maranon DG, Jimenez-Sainz J, Fontenay GV, Kwon Y, Leung SG, Lu L, Jensen RB, et al (2015) Promotion of BRCA2-Dependent Homologous Recombination by DSS1 via RPA Targeting and DNA Mimicry. Molecular Cell 59: 176–187

